# Stepped Cyclic Strain, that Increases or Decreases as Hierarchical Collagen Fibers Form, Does not Further Improve Maturation in Engineered Ligaments

**DOI:** 10.64898/2026.08.04.742835

**Authors:** Leia Troop, Jennifer L. Puetzer

**Affiliations:** Department of Biomedical Engineering, Virginia Commonwealth University, Richmond, VA, 23284, United States; Department of Orthopaedic Surgery, Virginia Commonwealth University, Richmond, VA, 23284, United States

## Abstract

The primary source of strength in ligaments and tendons are hierarchically organized collagen fibers. These fibers largely do not regenerate after injury, with repair, nor in engineered replacements, limiting treatment options. Previously, we developed a culture system which guides ACL fibroblasts in high-density collagen gels to form native-size hierarchical fibers over 6 weeks, and demonstrated that intermittent cyclic stretch further improves maturation. However, additional maturation is needed for clinical relevance. Interestingly, we found cyclic load affected cells differentially depending on the degree of organization, with 10% cyclic strain driving early improvements in unorganized gels and 5% strain being more beneficial later in culture once cells were on aligned fibers. Here, we explored whether a stepped cyclic load, that increased or decreased in strain magnitude as collagen fibers developed, further improved maturation. We hypothesized that progressively decreasing cyclic strain as organization increases would drive cells to produce more mature hierarchical fibers, resulting in stronger replacements. Controls had intermittent cyclic stretch at 0, 5, 7, or 10% strain throughout culture, while stepped load constructs were cyclically loaded with a strain that increased or decreased by 2-3% every 2 weeks as constructs matured. Contrary to our hypothesis, neither decreasing nor increasing load led to further tissue maturation. We hypothesize stepped cyclic load may disrupt cellular tensional homeostasis, leading to repeated remodeling of collagen and shifted proteoglycan accumulation. This study provides insight into how stepped cyclic loading affects hierarchical fiber formation and maturation, which will help to engineer stronger replacements and better rehabilitation protocols.

## 1. Introduction

Movement and stability within the musculoskeletal system is largely provided by tendons and ligaments which attach muscle-to-bone and bone-to-bone, respectively [1–4]. These tissues are composed of hierarchically organized collagen fibers that run the length of the tissue, consisting of aligned nanometer-wide fibrils that group together into larger fibers and fascicles [5,6]. This organization is critical to the function of tendons and ligaments, with each level reinforcing the overall fiber and providing the strength to withstand the repeated loads that occur in daily life [3,5–7]. In addition to the hierarchically organized collagen, multiple other components also help to further reinforce the strength of these fibers, including lysyl oxidase (LOX) crosslinks, proteoglycans, and crimp [3,8,9]. Altogether, these components provide tendons and ligaments with strong, non-linear tensile properties that are essential to long-term function [3,5–8,10,11]. The importance of this hierarchical organization to the function of tendons and ligaments is established, but recreating these fibers after injury or in engineered tissue replacements remains a challenge [12,13].

Injuries disrupt this collagen organization and reduce tendon and ligament function. Further, after these injuries, the collagen fibers largely do not regenerate and instead often heal with unorganized scar tissue, which reduces tissue function long-term [1,13,14]. Current repair options are limited to autograph or allograph replacement for large tears [4,15], or specialized repair techniques for partial tears [16–18], all of which have complications and high re-rupture rates [13,14,19–21]. Engineered replacements are a promising alternative, however it remains a challenge to regenerate the larger hierarchical fibers and fascicles of native tissue critical to meet long-term mechanical demands of daily loading [13,15,22–24]. Previously, we developed a novel culture system that uses static boundary restraints to guide Anterior Cruciate Ligament (ACL) fibroblasts and flexor tenocytes in high-density collagen gels to form hierarchical fibers that match immature tissue by 6 weeks of culture [25,26]. These constructs are promising, but further improvements are needed to be clinically relevant for adults.

It has been well established that mechanical cues are critical for tissue development *in vivo* [1,8,27] and have been shown to drive maturation in engineered tissues *in vitro* [28–39]. In particular, a large amount of work has explored the potential of cyclic loading, mirroring cyclic muscle activity. Cyclic load has largely been explored via 2D equiaxial strain [30,40–43], 2D or 3D uniaxial tension [27,33,39,44], and 3D hydrogels under multimodal loading such as tension and torsion simultaneously [30,43,45,46]. These studies have generally found that intermittent cyclic load [30,46,47], particularly at strains 1-5%, or within *in vivo* strains [43,45,46], improves fibril alignment, fibril diameter, and collagen production in engineered tendons and ligaments [1,8,27,28,30–32,34–36,39,48]. However, these studies have largely focused on cell differentiation alone [49] or evaluating these effects when collagen is organized to the nanometer fibril level only, and the effect of these loads as hierarchical fiber organization progresses to larger fibers and fascicles is largely unknown.

Recently, we evaluated the effect of intermittent cyclic load at 5 and 10% strain in our culture system which supports hierarchical fiber formation [39]. We found that both 5 and 10% strain drove increased fiber formation and tissue maturation, however cells responded differentially depending on the degree of organization [39]. More specifically, we found 10% cyclic strain drove early improvements in mechanics and composition when cells were in unorganized gels, and 5% load was more beneficial later in culture once cells were anchored on aligned collagen fibers, suggesting a shift in mechanotransduction [39]. It has been reported cells sense load more intensely when attached to aligned fibers [50], therefore it makes sense that a higher strain magnitude may be optimal when cells are in unorganized gels, but it may need to decrease as collagen fibers form so to not overload cells and trigger an injurious response. These results indicate that a loading regime that decreases in intensity from a higher strain early in culture to a lower strain later in culture may drive synergistic improvements or enhanced maturation.

However, adaptive loading regimes, which change with time in culture have had limited exploration in engineered tissues. Extensive work *in vivo* has evaluated the strain threshold for tendon and ligament remodeling and repair [51–54]. Tendons and ligaments respond, in the short-term and long-term, to load through remodeling, which changes the applied strain needed to achieve the same load. These studies have found that during healing, as collagen fibers reform, tissues require an increase in strain to maintain the same load and to continue to drive healing [55–58]. Further, in tendon and ligament development, cyclic load increases in strain magnitude during maturation [8,11]. Collectively, this suggests that rather than a decreasing strain, an increasing strain may be necessary to drive further maturation in our engineered tissues.

There remains a knowledge gap in how adaptive, stepped, or ramped loading regimes, which change magnitude, duration, or frequency as tissues develop, affect engineered tissues. Our previous data [39] and natural development [1,8] suggest a stepped load that changes in strain magnitude as cell organize the tissue can drive enhanced maturation over steady strain magnitudes. Therefore, the objective of this study was to explore whether a stepped cyclic load that changes intensity, by either increasing or decreasing by 2-3% strain as collagen fibers develop, further improves engineered ligament maturation. We hypothesize that decreasing cyclic strain as fiber organization increases will drive cells to increase tissue maturation, producing significantly stronger replacements. A better understanding of how cells respond to applied load at different levels of collagen organization could not only help to produce more mature, clinically relevant engineered replacements but also help inform loading protocols across different engineered systems, and help develop more optimal rehabilitation protocols.

## 2. Methods and Materials

### 2.1 Cell isolation and construct fabrication

Ligament fibroblasts were isolated from neonatal bovine as previously described [25,26,39,48]. Briefly, 1–3-day old neonatal bovine legs were purchased from a slaughterhouse, and cells were isolated within 48 hours of culling. The bovine cranial cruciate ligament (CCL), the analog to human ACL which will thus be referred to as bovine ACL, was aseptically isolated, diced, and enzymatically digested in 0.2% collagenase. Cells from 3 separate bovine were combined to limit donor variability, passaged once to obtain enough cells for construct formation, and then seeded into high-density collagen gels. Native ACL samples, used for imaging and composition analysis, were collected at the time of isolation and stored as described in sections 2.4 Hierarchical Collagen Organization Analysis and 2.6 Compositional Analysis.

High-density, cell-laden collagen gels were fabricated as previously described [25,26,39,48,59]. Briefly, tendons from equal numbers of male and female Sprague-Dawley rat tails (BIOVT) were aseptically removed and stored in 0.1% acetic acid to isolate collagen from the tendons. The collagen was then frozen, freeze-dried, and reconstituted to 30 mg/ml collagen in 0.1% acetic acid to form stock solutions [25,60]. To form constructs, the stock collagen solution was mixed with a working solution of 1x PBS, 10x PBS, and 1 M NaOH to neutralize the solution and initiate gelation [25]. Ligament fibroblasts were immediately mixed into the collagen solution to ensure cells were seeded evenly throughout the gel. The collagen-cell solution was cast into 1.5 mm thick sheet gels at 20 mg/mL collagen and 5 million cells/mL [25,26,39,48,59]. The gels were set at 37°C for one hour and then cut into individual constructs with an 8x30 mm die, yielding 4-6 constructs per sheet gel. Constructs were divided between groups and cultured for up to 6 weeks. To make each sheet gel, different collagen stocks and cell isolations were used. Thus, N refers to individual constructs from different collagen stocks and cell expansions.

### 2.2 Culture Conditions and Mechanical Stimulation

Constructs were either cultured static in our clamping device, or clamped in a tensile bioreactor and loaded with steady intermittent cyclic load that remained at the same strain throughout culture or with a stepped strain that changed as aligned fibril, fibers, and fascicles formed. More specifically, constructs were clamped into their designated culture device 24 hours after generation as previously described [25,26,39,39,59]. Static constructs (0% load) were cultured in our static clamping device (**Figure 1A**) which we have previously demonstrated guides cells to produce hierarchical fibers over 6 weeks [25,26,59], while loaded constructs were clamped into a modified CellScale tensile bioreactor (**Figure 1B**) [39,48], with both setups having a 20 mm gauge length between clamps. Loaded constructs were stimulated with an established loading regime we have shown previously drives maturation in our system [39,48], consisting of intermittent cyclic stretch at 1 Hz, for 1 hour on, 1 hour off, 1 hour on, 3 times per week on Monday, Wednesday, and Friday (**Figure 1C**). Steady load control constructs were loaded at 5, 7, or 10% cyclic strain throughout culture. The control 7% strain data is not included in the manuscript figures to allow for easier interpretation of results, but are provided in supplemental (**Supplemental Figures 1-4**). Stepped load constructs were loaded with a strain that changed as constructs matured. Specifically, increasing load constructs were loaded at 5% strain from weeks 0-2, 7% strain weeks 2-4 once aligned fibrils formed and 10% strain weeks 4-6 once aligned fibers formed, while decreasing load constructs were loaded with the opposite pattern (**Figure 1D**). All constructs were cultured in a standard media composed of Dulbecco’s Modified Eagle Medium (DMEM) with 10% fetal bovine serum (FBS), 1% antibiotic/antimycotic, 50 µg/ml ascorbic acid, and 0.8 mM L-proline [25,39,48]. Conditional media changes were performed every 2-3 days prior to loading, replacing half of the media during each media change [48].

**Figure 1.**
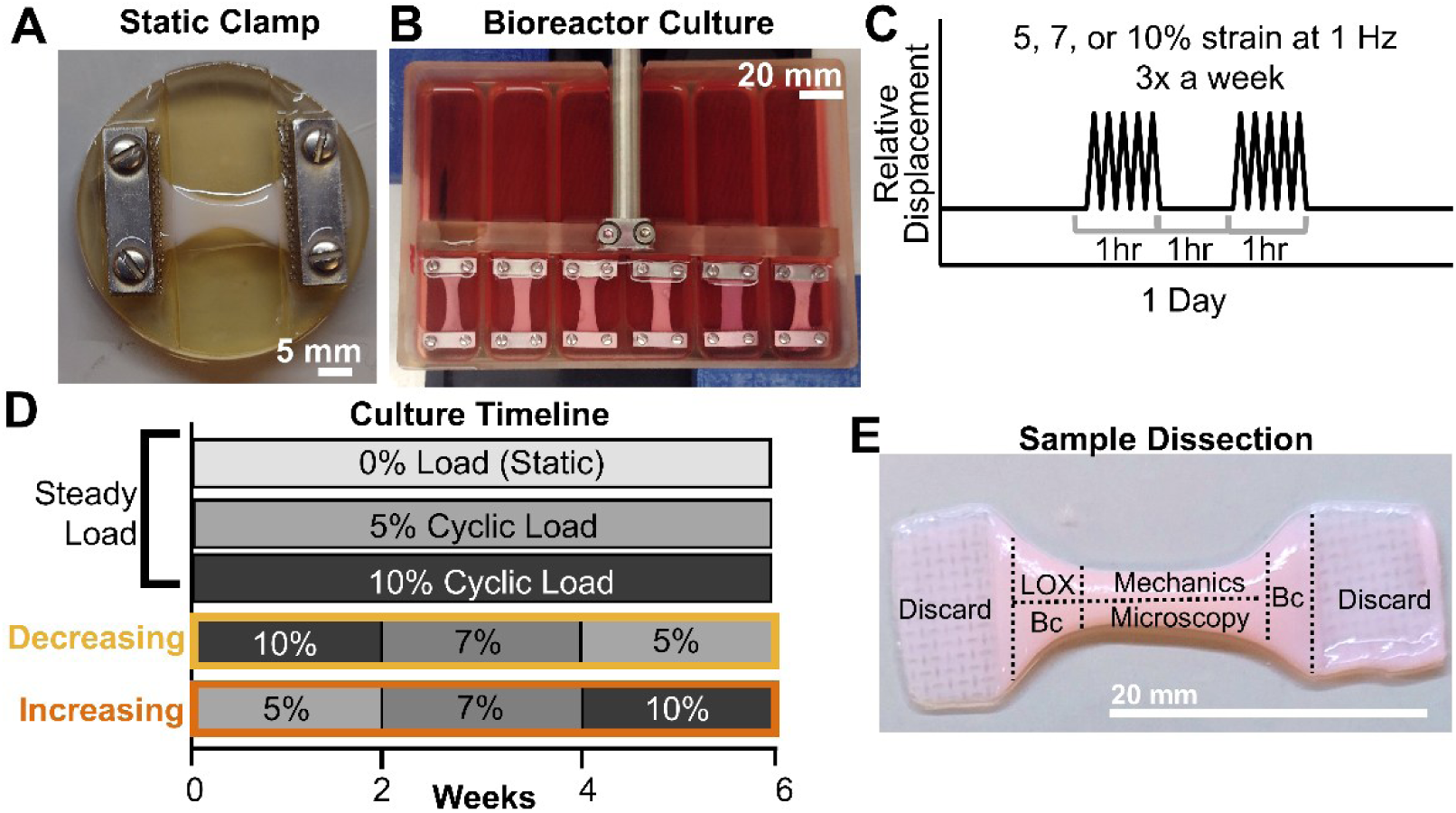
Experimental setup with constructs either statically clamped or cyclically loaded for up to 6 weeks. A) Static clamping culture device and B) bioreactor setup for cyclically loaded constructs. C) Depiction of intermittent loading regime. Constructs were loaded with an established loading regime, 3 times a week (MWF) for 1 hour on, 1 hour off, 1 hour on at 1 Hz and 5%, 7% or 10% strain. D) Control groups were kept statically clamped or loaded with steady intermittent cyclic loading at 5% or 10% strain throughout culture, while experimental stepped load groups were loaded with a cyclic strain that decreased or increased as organization improved at 2 and 4 weeks of culture. E) Depiction of tissue sectioning performed at each timepoint. Clamped regions were removed and remaining tissue was allocated for composition (Bc), lysyl oxidase activity (LOX), mechanical, and microscopy analysis, including collagen organization analysis and immunohistochemistry analysis of proteoglycans.

### 2.3 Post-culture Collection and Analysis

Timepoints were collected at 0, 2, 4, and 6 weeks, with 0-week timepoints collected 24 hours after clamping and/or one loading cycle. For each timepoint, 6-10 constructs per group were collected, weighed, photographed, and sectioned to analyze collagen organization, tissue tensile properties, composition, LOX activity, and proteoglycan localization (**Figure 1E**). Due to significant contraction, not all sections could be taken from all constructs, therefore later timepoints of loaded groups have a larger number of constructs cultured than static controls. Upon removal from culture, constructs were photographed and weighed to determine contraction over time in culture. Percent original area was determined using FIJI (NIH) and normalized to respective 0 week constructs as previously described [26,48]. Construct percent weight was determined by measuring whole construct weight at each time point and comparing to respective 0 week constructs.

### 2.4 Hierarchical Collagen Organization Analysis

Assessment of hierarchical fiber formation was performed at the fiber-scale (1-100 µm) via confocal microscopy, at the fascicle-scale (>100 µm) via picrosirius red staining, and at the fibril scale (<1 µm) via scanning electron microscopy (SEM) in constructs and native tissues, as previously reported [26,39,48,59]. Briefly, samples spanning the length of the construct (**Figure 1E**) or native ACL were fixed in 10% formalin and stored in 70% ethanol. A total of 6-8 constructs per timepoint and treatment, and 4 native ACLs were imaged and analyzed via confocal reflectance. After confocal imaging, 6-week samples from each treatment were split into groups for picrosirius red staining (N = 3) or SEM analysis (N = 3).

#### 2.4.1 Confocal Reflectance Imaging and Analysis

Confocal reflectance imaging for fiber-level analysis was performed with a Zeiss LSM 980 microscope and Plan-Apochromat 20x/1.2 objective as previously described [25,38,39,61]. Briefly, collagen was visualized with a 405 nm laser by capturing reflectance through a 27 µm pinhole at 400-465 nm. At the same time, auto-fluorescence of cells was captured with a 488 nm laser at a range of 509-571 nm. Confocal images taken across the constructs (N = 6-8 constructs, n = 6-8 images per construct) and neonatal bovine ACLs (N = 4 samples, n = 6-8 images per sample) were analyzed using a custom Fast Fourier transform (FFT) based MATLAB code as previously described [39,62]. Briefly, the degree of collagen alignment was determined using an established alignment index measure based on the directionality of the FFT, where 1 is unorganized and 4.5 is completely aligned [62]. Images were then rotated to the major axis of alignment, a second FFT was performed, and the mean fiber diameter within the image was determined from the average frequency of the FFT [39,62]. Values from representative images for each construct were pooled to determine average alignment and diameter for each construct (n = 6-8 images per construct). The reported alignment and diameter values are the average of construct values (N = 6–8 for constructs, N =4 for neonatal ACL).

#### 2.4.2 Polarized Picrosirius Red Imaging

Polarized picrosirius red imaging was performed to observe fascicle-level (>100 µm scale) organization and crimp as previously described [25,26,39,48,59]. Briefly, 6-week constructs (N = 3) and neonatal bovine ACL (N = 3) were embedded in paraffin, sectioned, and stained with picrosirius red. Slides were then imaged with a Nikon Eclipse Ts2R inverted microscope and Nikon Pan Fluor 10x/.30 OFN25 Ph1 DLL objective in linear polarized light.

#### 2.4.3 SEM Imaging and Analysis

SEM imaging to analyze fibril-level (<1 µm scale) organization was conducted on 6 week constructs and neonatal bovine ACL with a Hitachi SU-70 FE-SEM as previously described [26,48,59]. Briefly, fixed samples were frozen and 10-12 slices, 5 µm thick, were removed from the surface using a Leica CM 1860 UV cryostat to expose fibrils within constructs and create a flat surface for imaging. Samples were then washed in PBS, serially shifted to 100% ethanol, dried via critical point drying, mounted cut side up on 1 inch aluminum mounts, coated with 0.025–0.035 kAngstroms platinum, and imaged at working distance 10 mm, with 5 kV, at 10,000X and 50,000x magnification. Images at 50,000X were analyzed to determine fibril alignment and diameter as previously described [26,39]. Briefly, alignment was measured via the FIJI directionality function to determine degrees of dispersion, with 6-8 images per sample averaged to determine construct fibril dispersion. Bar graphs are the average of construct dispersion values (N = 3). Fibril diameter was determined by using the FIJI measure function to record the diameter of 20 fibrils per image that were pooled (total 120 fibrils per construct or ACL) to determine average fibril diameter for each construct [39,59]. All measured diameters from N = 3 samples were pooled (n = 360) for violin plots to visualize spread of diameters at 6 weeks. Construct values were statistically analyzed by averaging 120 fibril diameters per sample to compare construct fibril diameter averages at 6 weeks (N = 3 for constructs and native ACL).

### 2.5 Mechanical Analysis

Tensile tests were performed as previously described with a BOSE ElectroForce 3200 System equipped with a 250 g load cell and serrated grips [25,26,39,48,59]. Briefly, lengthwise samples (N = 6-8) were collected from each construct and stored at −20°C. Prior to testing, samples were thawed in PBS with EDTA-free protease inhibitor, measured, and stretched to failure at 0.75% strain/second, assuming quasi-static loading, as with previous studies [25,39,59]. Tensile properties were determined via a previously developed linear regression-based MATLAB code [48]. Briefly, the toe and elastic moduli were determined by fitting the two regions with individual linear regressions, where r^2^ > 0.999. Toe transition stress and strain were defined as the point where the 2 fit lines intersect, and ultimate tensile strength (UTS) and failure strain were defined as the point of maximum stress and strain before failure, respectively.

### 2.6 Compositional Analysis

DNA, collagen, glycosaminoglycan (GAG), and lysyl oxidase (LOX) activity analysis were performed as previously described for constructs and native samples [25,26,39,48,59]. Briefly, samples collected for DNA, collagen, and GAG analysis (N = 6-8, **Figure 1E**) were weighed wet, frozen, lyophilized, and weighed dry before digestion in 1.25 mg/ml papain solution at 60°C for 16 hours. DNA, collagen, and GAG concentrations were determined via a modified Quant-iT PicoGreen dsDNA assay kit (Invitrogen), a modified hydroxyproline (hypro) assay [63], and a 1,9- dimethylmethylene blue (DMMB) assay at pH 1.5 [64]. Reported values are normalized to sample dry weight (DW). Samples collected for LOX activity analysis (N = 5-7, **Figure 1E**) were stored in a 6 M Urea 10mM Tris-HCl solution at pH 7.4 with 1% protease inhibitor at each time point and frozen at −80°C as previously described [39,59]. Arbitrary units (A.U.) of LOX activity were determined via a fluorometric LOX activity assay (Abcam, ab112139). Results were normalized to sample DNA measured via Quant-iT Picogreen dsDNA assay.

### 2.7 Proteoglycan Localization

Based on the significant increase in GAG accumulation at 6 weeks in stepped load constructs, all cultures were evaluated for small leucine rich proteoglycan (SLRP) and aggrecan localization at 6 weeks to better understand which types of proteoglycans were being accumulated. Static clamped constructs at 0 weeks and native ACL were also evaluated for comparison. Proteoglycan localization was evaluated using an established immunofluorescent protocol [26,48]. Briefly, representative 0-week 0% load constructs, 6-week constructs from each condition, and neonatal ACLs were fixed, embedded in paraffin blocks, and sectioned (N = 3). Sections were treated with proteinase K to retrieve antigens, blocked with 5% goat serum, and incubated overnight with primary polyclonal rabbit antibodies in 5% goat serum at dilution 1:200 for decorin (Kerafast LF- 94), biglycan (Kerafast LF-96), and lumican (Novus NBP1-87726) and at dilution 1:150 for fibromodulin (Fmod, Genetex-GTX54035) and aggrecan (Genetex-GTX45920). Negative controls were incubated overnight in 5% goat serum. All sections were then incubated with goat-anti-rabbit IgG secondary antibody labeled with Alexa Fluor 488 (Invitrogen A11008) at 1:200 dilution for 2 hours and cross-labeled with DAPI at 1:1000 dilution. Sections were imaged on Zeiss LSM 980 microscope with a Plan-Apochromat 40x/1.1 water objective, using the same reflectance and fluorescence settings as described in section 2.4.1 Confocal Reflectance Imaging and Analysis to capture both the collagen organization using confocal reflectance and the immunofluorescence simultaneously.

For analysis of proteoglycan localization, 6 representative images were taken from each section (N = 3 constructs per treatment). Images were analyzed in FIJI for percent area positive stain for each 353.55 x 353.55 µm (1024x1024 pixel) image. Negative controls were imaged with the same settings as positive samples, ensuring little-to-no signal was present as noise (**Supplemental Figure 5**).

### 2.8 Statistics

All data were analyzed with Shapiro-wilk tests to confirm normality within groups. After confirming normality, percent original area and weight, fiber alignment index and diameter, tensile properties, composition, and proteoglycan accumulation were analyzed via 2-way ANOVA with Tukey’s post-hoc, with p < 0.05 as significant (SigmaPlot 14). For image and compositional analysis, where multiple measurements per construct were taken, measurements from each construct were pooled and averaged to determine an overall value for each construct. Statistical analysis was performed using these construct averages. Fibril dispersion and fibril diameter data were only collected at 6 weeks, thus 1-way ANOVA was used for analysis, with the investigated effect only being culture condition. All data are expressed as mean ± standard error (S.E.M.).

## 3. Results

### 3.1 Tissue Gross Morphology

Inspection of construct gross morphology revealed that all constructs contracted over 6 weeks of culture, with steady cyclic load constructs appearing to have greater contraction than static and stepped load constructs (**Figure 2A**). This observation was supported by percent area measurements, with steady load constructs having significantly smaller area than static constructs at 4 and 6 weeks (**Figure 2B**). Stepped load constructs generally did not contract to the same degree as steady loaded constructs, with increasing load constructs having significantly greater percent area compared to steady load by 6 weeks. However, despite differences in percent area, all groups had similar percent weight by 6 weeks, except for decreasing load constructs which had a significantly greater wet weight than all other loaded groups at 4 and 6 weeks, potentially indicating increased swelling.

**Figure 2.**
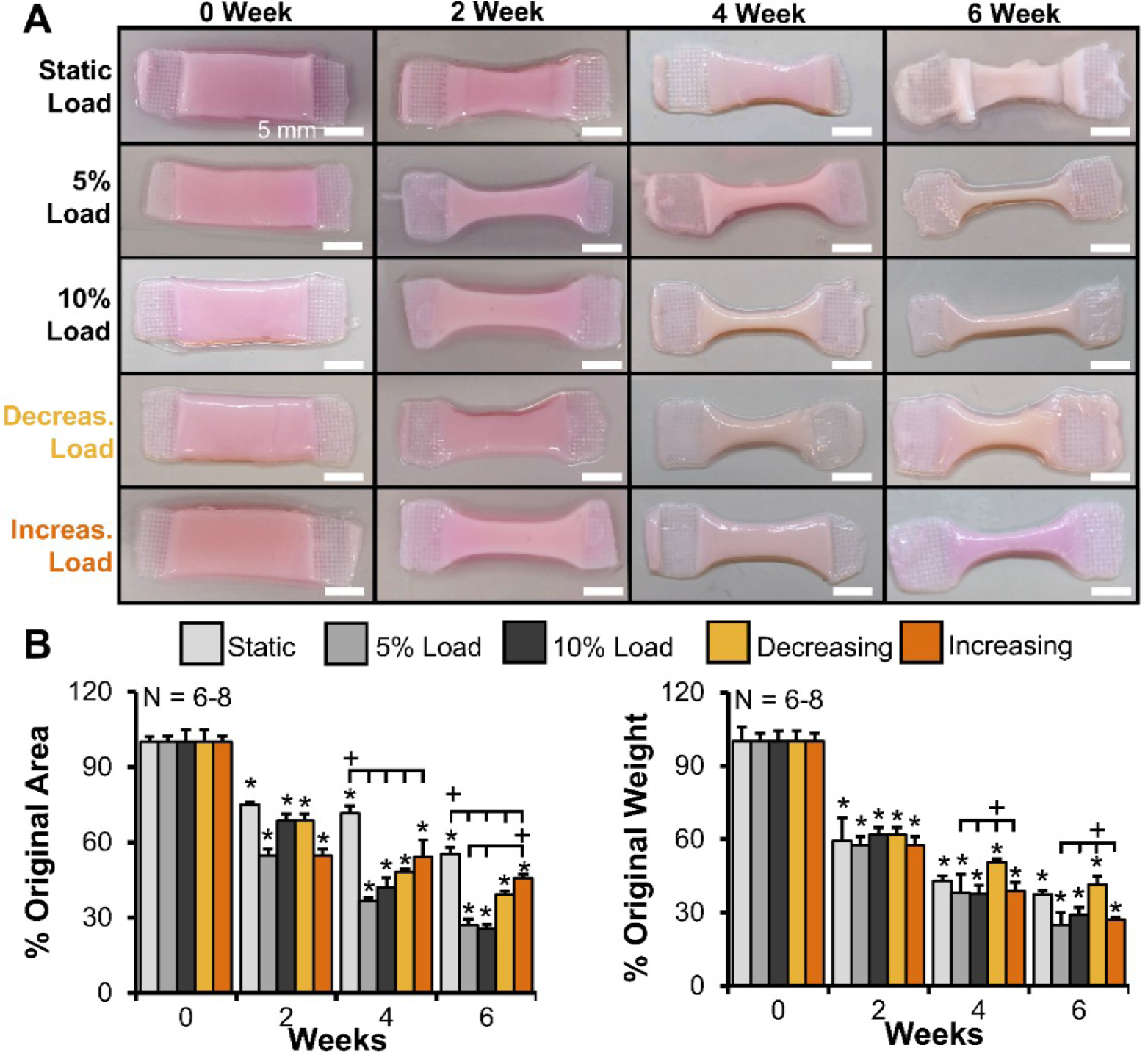
Stepped load constructs appeared to have reduced contraction compared to steady load constructs with time in culture. A) Photographs of representative constructs at each time point (Scale bar = 5 mm). B) Percent area and percent wet weight throughout culture compared to respective 0 week constructs (N = 6-8). Data shown as mean ± S.E.M. Significance compared to *0 week static or +bracket group (p < 0.05).

### 3.2 Tissue Hierarchical Collagen Organization

Confocal reflectance imaging at the fiber-scale revealed all culture conditions guided cells in unorganized collagen at 0 weeks to form aligned collagen fibrils by 2 weeks, and larger fibers and fascicles by 4 and 6 weeks of culture, similar to previous studies (**Figure 3A**) [25,26,39,59]. Further, steady load drove enhanced fiber development, with collagen fibers appearing larger and more organized than static constructs at 4 and 6 weeks, similar to our previous study [39]. However, both increasing and decreasing stepped load constructs appeared to have less collagen crimp and reduced organization compared to steady load constructs, with organization more similar to static controls at 4 and 6 weeks.

**Figure 3.**
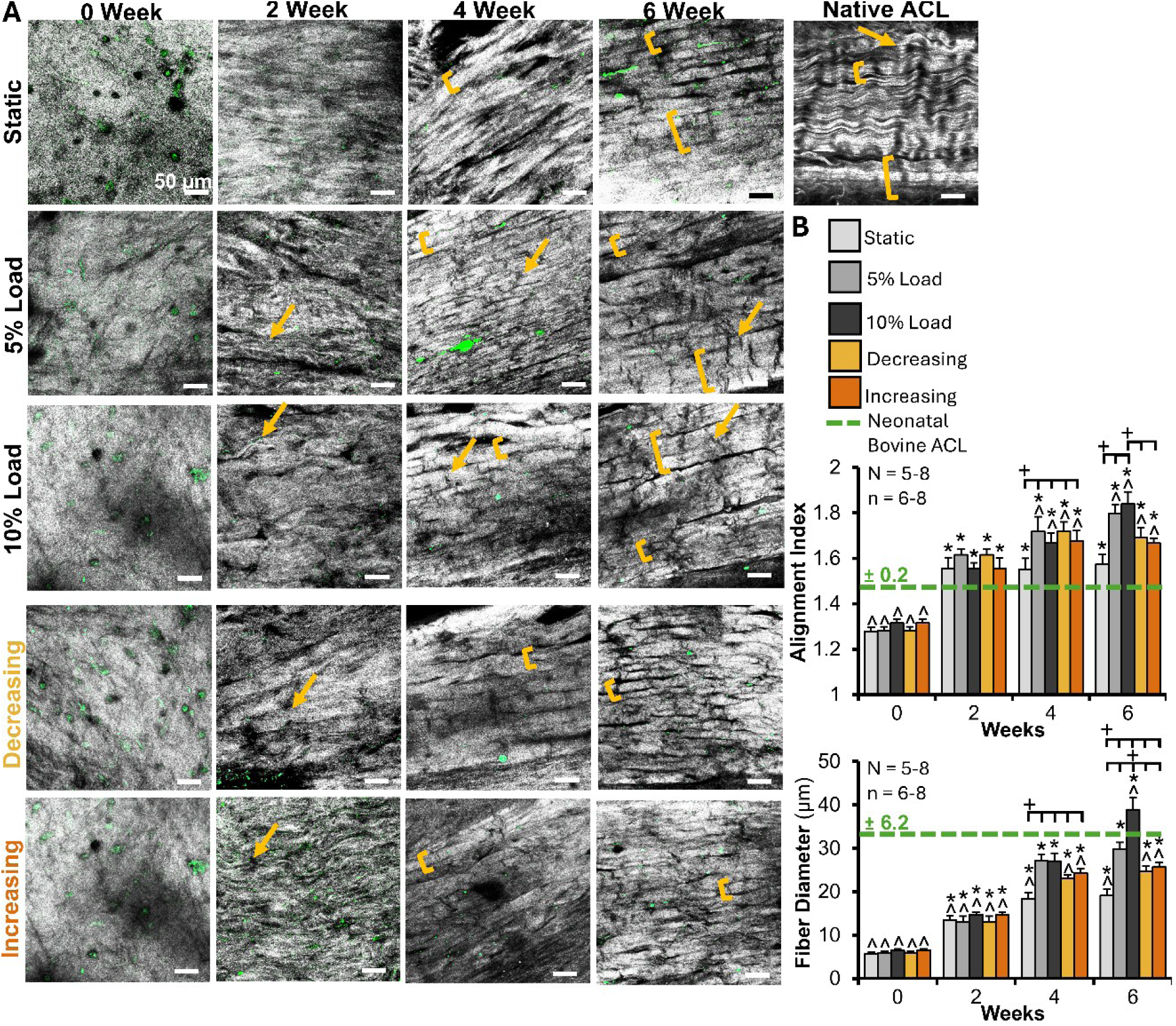
Stepped load constructs had reduced fiber development compared to steady load constructs. A) Confocal reflectance revealed all groups develop aligned fibrils by 2 weeks, and larger fibers by 4 weeks (brackets), with steady loading producing increased fiber and early fascicle formation (larger brackets), and increased crimp formation (arrows) compared to static and stepped load constructs. Grey = collagen, green = cellular auto-fluorescence. Scale bar = 50 µm. B) Degree of collagen alignment (reported via alignment index where 1 is unorganized, 4.5 is perfectly aligned) and average collagen fiber diameter determined via a FFT-based image analysis. 6–8 images per construct were averaged to determine construct values (N = 6-8 constructs per time point and N = 4 neonatal bovine ACL). Data shown as mean ± S.E.M. Significant difference compared to *0 week static, ^neonatal bovine ACL, or +bracket group (p < 0.05).

Analysis of confocal images revealed that all groups had significant improvements in alignment and fiber diameter with time in culture, and reached native levels of alignment by 2 weeks (**Figure 3B**). Steady load drove further improvements, with constructs having significantly increased alignment and fiber diameters over static constructs at 4 and 6 weeks, and average fiber diameters that were not significantly different from neonatal bovine ACLs by 4 weeks. Steady 10% load produced the largest fiber diameters by 6 weeks, surpassing native tissue and all other groups. Stepped load constructs also continued to mature and develop more aligned fibers through 6 weeks of culture, but not to the same extent as steady load. By 6 weeks, stepped load constructs had greater alignment than neonatal bovine ACL, but not static constructs. Further, stepped load constructs had significantly larger fiber diameters than static constructs at 4 and 6 weeks, but did not reach native or 10% load fiber diameter.

Polarized picrosirius red analysis of the fiber- and fascicle-length scales at 6 weeks further confirmed that increasing and decreasing stepped load, which changed as hierarchical fibers formed, did not improve fiber maturation compared to steady load. Low magnification images (10x) demonstrated that while steady load constructs developed early fascicles, stepped load constructs did not experience further bundling and appeared to have far less organization overall (**Figure 4A**). Higher level magnification (30x images) revealed that steady load constructs developed fibers with large, more defined, and regularly spaced crimps, while stepped load constructs appeared to have disrupted crimps that were smaller, less defined, and less regularly spaced.

**Figure 4.**
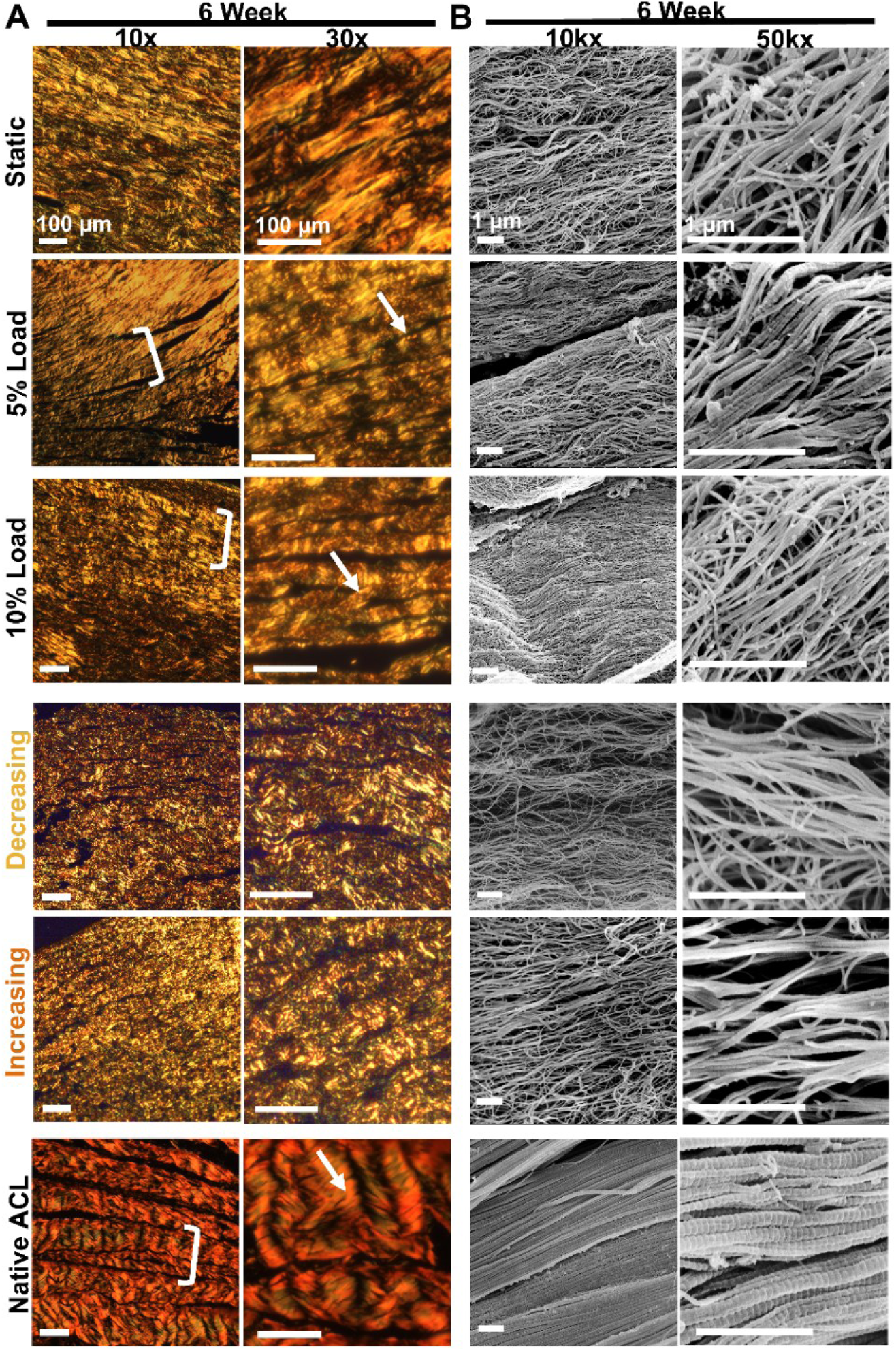
Stepped load constructs had reduced fascicle and fibril organization at 6 weeks compared to steady load constructs. A) Fascicle length-scale organization at 6 weeks evaluated by picrosirius red staining, imaged with polarized light, (scale bar = 100 µm). Steady loading drove enhanced fascicle formation and more regular crimp formation (arrows) compared to static and stepped load constructs. B) SEM images of 6-week constructs to assess fibril length-scale organization (scale bar = 1 µm). Steady load constructs had clear bundling of fibrils into larger fibers by 6 weeks, while this was largely absent from stepped load constructs.

Similar to fiber-level organization, SEM imaging revealed that all constructs formed aligned fibrils by 6 weeks (**Figure 4B**). Further, steady load appeared to drive increased maturation with fibrils condensing into larger fibers, similar to native tissue. Conversely, stepped load constructs largely lacked these groupings of fibrils. Analysis of fibril-level dispersion revealed that steady load significantly reduced fibril dispersion compared to static clamping, representing an increase in fibril alignment (**Figure 5**). By 6 weeks, both 5% and 10% steady load constructs were not significantly different from native levels of fibril dispersion. Conversely, both stepped load constructs had significantly greater dispersion than native tissue, i.e. less alignment, and decreasing load constructs had greater dispersion than 10% steady load at 6 weeks. Fibril diameter analysis indicated that all loaded constructs formed fibrils with larger diameters than static clamping alone, and 10% steady load produced average fibril diameters not significantly different from that of neonatal bovine ACL by 6 weeks of culture. Further, while both stepped load constructs developed more variable, larger fibril diameters than 5% steady load, both groups did not mature to match 10% steady load or neonatal bovine ACL.

**Figure 5.**
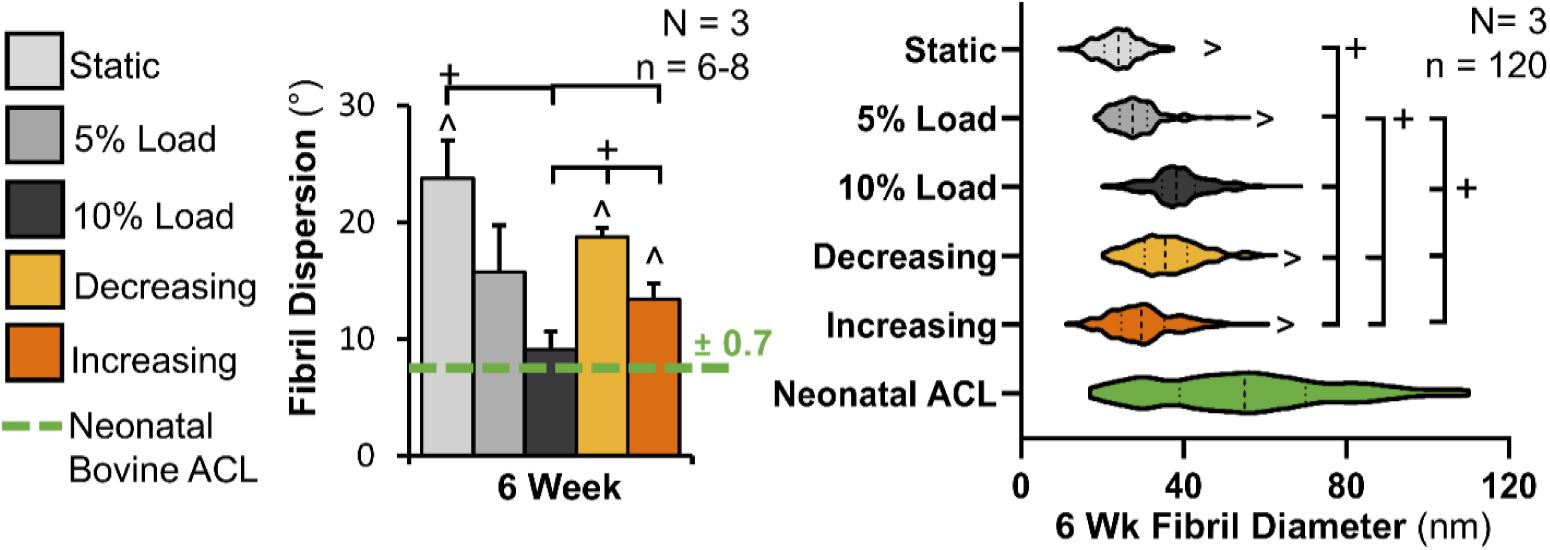
Stepped load had reduced fibril alignment and fibril diameters compared to 10% steady load constructs. Analysis of SEM images to determine fibril dispersion (lower dispersion indicating increased alignment), and fibril diameter of 6-week constructs and neonatal bovine ACL. For dispersion, 6–8 images per construct were averaged to determine the construct average (N = 3 for constructs and native ACL). For fibril diameters, 20 fibrils per image for 6 images were measured and pooled (total 120 fibrils per construct) to determine the average fibril diameter for each construct and these average construct values were used for statistical analysis. All fibril diameters from N = 3 constructs were pooled (n = 360) to visualize the spread of fibril diameters at 6 weeks. Data shown as mean ± S.E.M. Significance compared to ^neonatal bovine tissue or +bracket group (p < 0.05).

### 3.3 Tensile Mechanical Properties

Mirroring organization, all constructs had improved elastic tensile properties by 6 weeks (**Figure 6A**), with steady 5% load developing the highest elastic modulus and ultimate tensile strength (UTS) of all groups by 6 weeks, similar to our previous study [39]. While stepped load constructs did have significant improvements in tensile properties by 6 weeks, both decreasing and increasing load constructs had significantly lower elastic modulus and UTS than static and steady load constructs by 6 weeks. Similarly, steady load drove significant improvements in toe-region properties (**Figure 6B**), with both groups developing significantly greater toe modulus and transition stress compared to all other groups by 6 weeks. Stepped load constructs again had significant improvements in toe-region properties by 6 weeks, but these improvements were similar to static constructs and significantly less than steady load constructs. However, all loaded groups had similar significant improvements in failure strain and transition strain compared to static constructs at 6 weeks, decreasing toward native strain properties.

**Figure 6.**
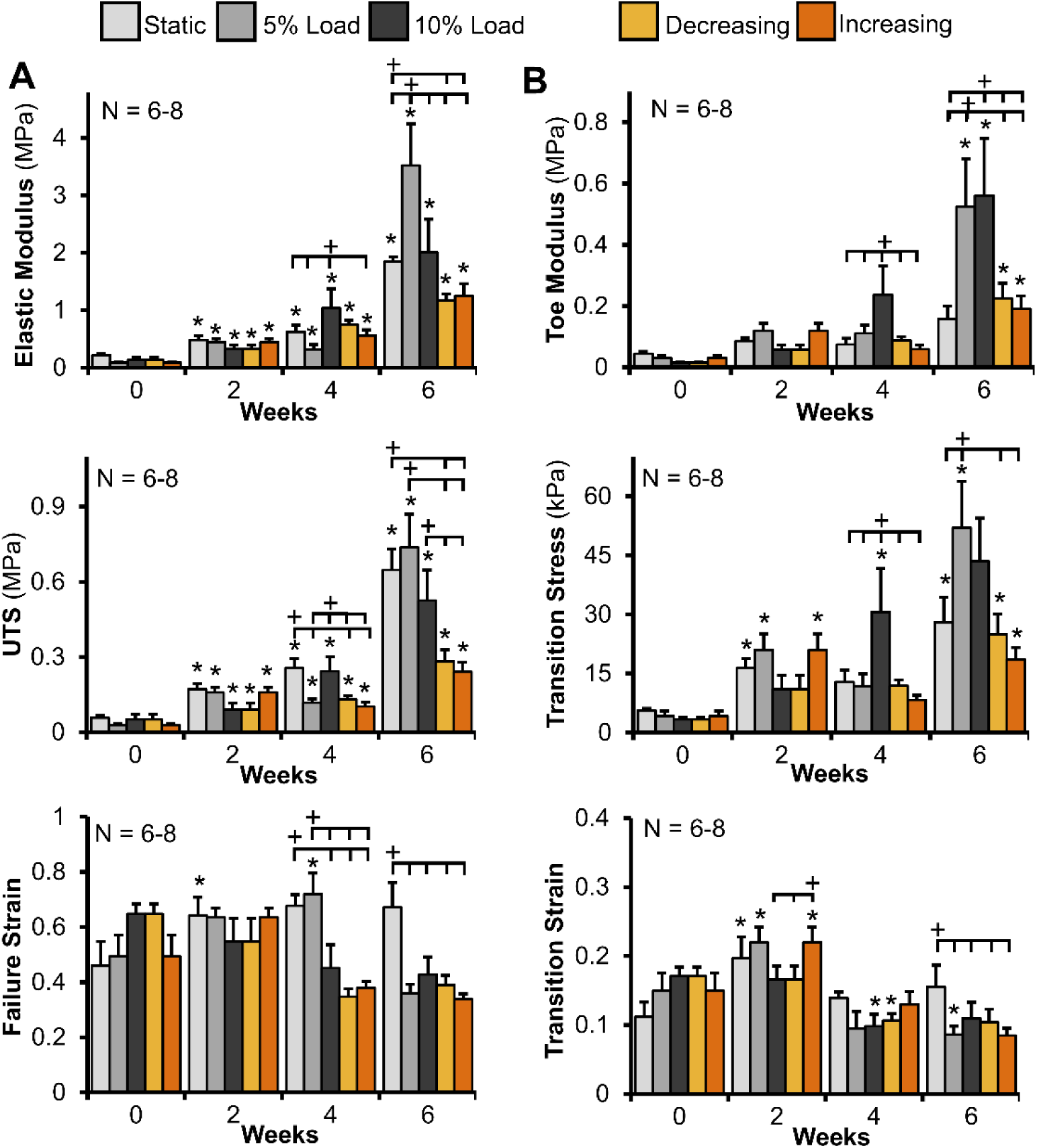
Stepped load did not improve tissue tensile properties over that of steady load constructs. Stepped load constructs had improved A) elastic properties (Elastic modulus, ultimate tensile strength (UTS), and failure strain) and B) toe-region properties (Toe modulus, transition stress, and transition strain) over 6 weeks of culture, but did not improve to the same degree as steady load constructs. N = 6–8. Data shown as mean ± S.E.M. Significance compared to *0 week static and +bracket group (p < 0.05).

### 3.4 Tissue Composition and LOX Activity

The effect of stepped load on composition varied with time in culture, degree of collagen organization, and percent applied strain. DNA accumulation was initially accelerated with load, with all loaded groups having significantly increased DNA compared to static constructs by 2 weeks (**Figure 7**). Steady load constructs then leveled off at native levels, while stepped load constructs were at or below native concentration at 4 and 6 weeks. Steady cyclic load drove a significant increase in collagen content, with 5% load constructs surpassing neonatal bovine ACL collagen concentration by 6 weeks. Stepped load constructs had relatively no changes in collagen concentration through 4 weeks. However, between 4 and 6 weeks, decreasing load constructs had an increase in collagen concentration after being subjected to 5% strain, while increasing load constructs had a decrease in collagen concentration after being subjected to 10% strain.

**Figure 7.**
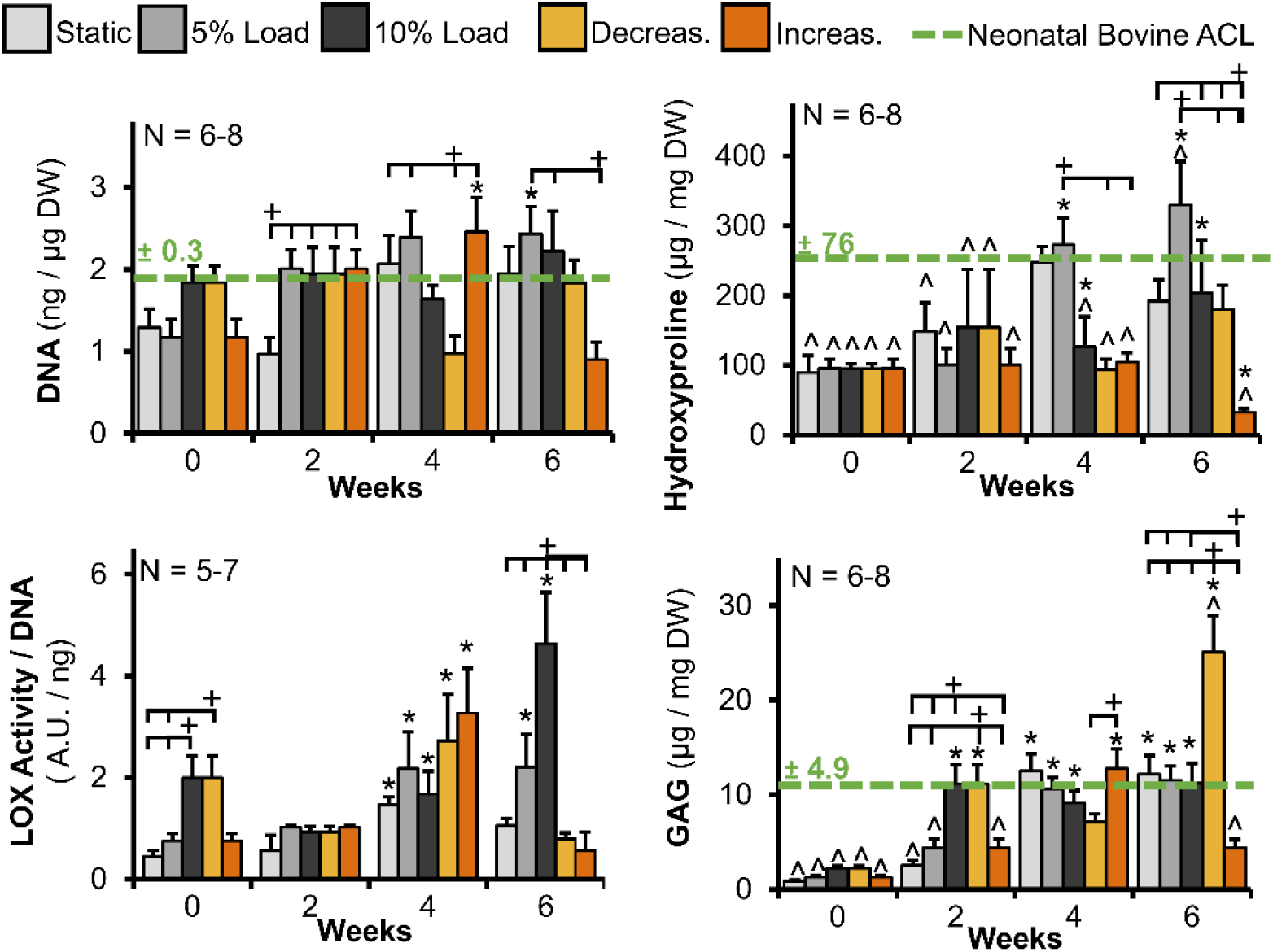
Stepped load constructs had reduced collagen, reduced LOX activity, and altered GAG accumulation compared to steady load constructs by 6 weeks. DNA, collagen content (represented by hydroxyproline), and GAG, normalized to dry weight (DW, N = 6-8), and LOX activity normalized to DNA (N = 5-7). Data shown as mean ± S.E.M. Significance compared to *0 week static, ^neonatal bovine ACL, and +bracket group (p < 0.05).

Looking at LOX activity, 10% load significantly increased LOX activity at 0 weeks, and all groups had significantly increased LOX activity at 4 weeks. However, by 6 weeks, only steady load groups had significantly increased LOX activity, with 10% steady load having the highest of all groups. Conversely, neither stepped decreasing nor increasing load had improved LOX activity at 6 weeks.

Finally, looking at GAG accumulation, steady 10% load drove accelerated GAG accumulation, reaching native levels by 2 weeks. Static and steady load constructs continued to accumulate GAG through 4 weeks of culture, leveling off at native concentrations. Conversely, decreasing load constructs continued to accumulate GAG, accumulating significantly higher GAG concentrations than all other groups, including neonatal bovine ACL, by 6 weeks. This significantly greater accumulation at 6 weeks is also present when normalizing to wet weight (**Supplemental Figure 6**), which when paired with the significantly higher percent weight of decreasing load constructs, indicates swelling in addition to GAG accumulation. Conversely, increasing load constructs reached native concentrations of GAG by 4 weeks, but then at 6 weeks, after experiencing 10% applied strain for 2 weeks, had a significant reduction in GAG content.

### 3.5 Proteoglycan Characterization and Localization

Due to the significant increase in GAG accumulation in decreasing load constructs, we were interested in evaluating which types of GAGs were accumulating. Tendons and ligaments accumulate two main classes of proteoglycans, composed of GAGs. These proteoglycan classes are small leucine rich proteoglycans (SLRPs) and large modular proteoglycans [65,66]. Generally, SLRPs, including biglycan, decorin, lumican, and fibromodulin, accumulate throughout hierarchical fiber development, modulating fibrillogenesis and fiber bundling [9,66–68]. Larger proteoglycans, like aggrecan, also accumulate during development, serving a role in fiber spacing, load distribution, and water retention [9,65,69,70]. Aggrecan is critical to tissue function, but excess of this proteoglycan in tendons and ligaments can indicate degeneration and injury [71,72]. Thus, we were interested in evaluating proteoglycan localization in our engineered ligament, and more specifically which class of GAGs was driving the increase in the decreasing load constructs.

Imaging revealed that all groups appeared to accumulate SLRPs and aggrecan with similar ratios to that of native tissue (**Figure 8**). In particular, all groups appeared to accumulate more decorin than all other SLRPs and to have increased aggrecan with time in culture, reaching staining intensities similar to native tissue. Major differences from native tissue were primarily in biglycan accumulation, with engineered tissues appearing to accumulate less biglycan than neonatal bovine ACL by 6 weeks. Further, stepped load constructs appeared to accumulate reduced biglycan compared to static and steady load controls, and instead had increased lumican accumulation compared to all other groups. Both stepped load groups appeared to have similar levels of fibromodulin, and higher levels of aggrecan than steady load constructs. Of the evaluated proteoglycans, most appeared to localize to the cell or between fiber bundles (**Figure 8**). This result supports proteoglycan roles in fiber spacing [68,69,73]. Decorin, however, appeared to have a more diffuse presence, covering entire fiber bundles, similar to native tissue.

**Figure 8.**
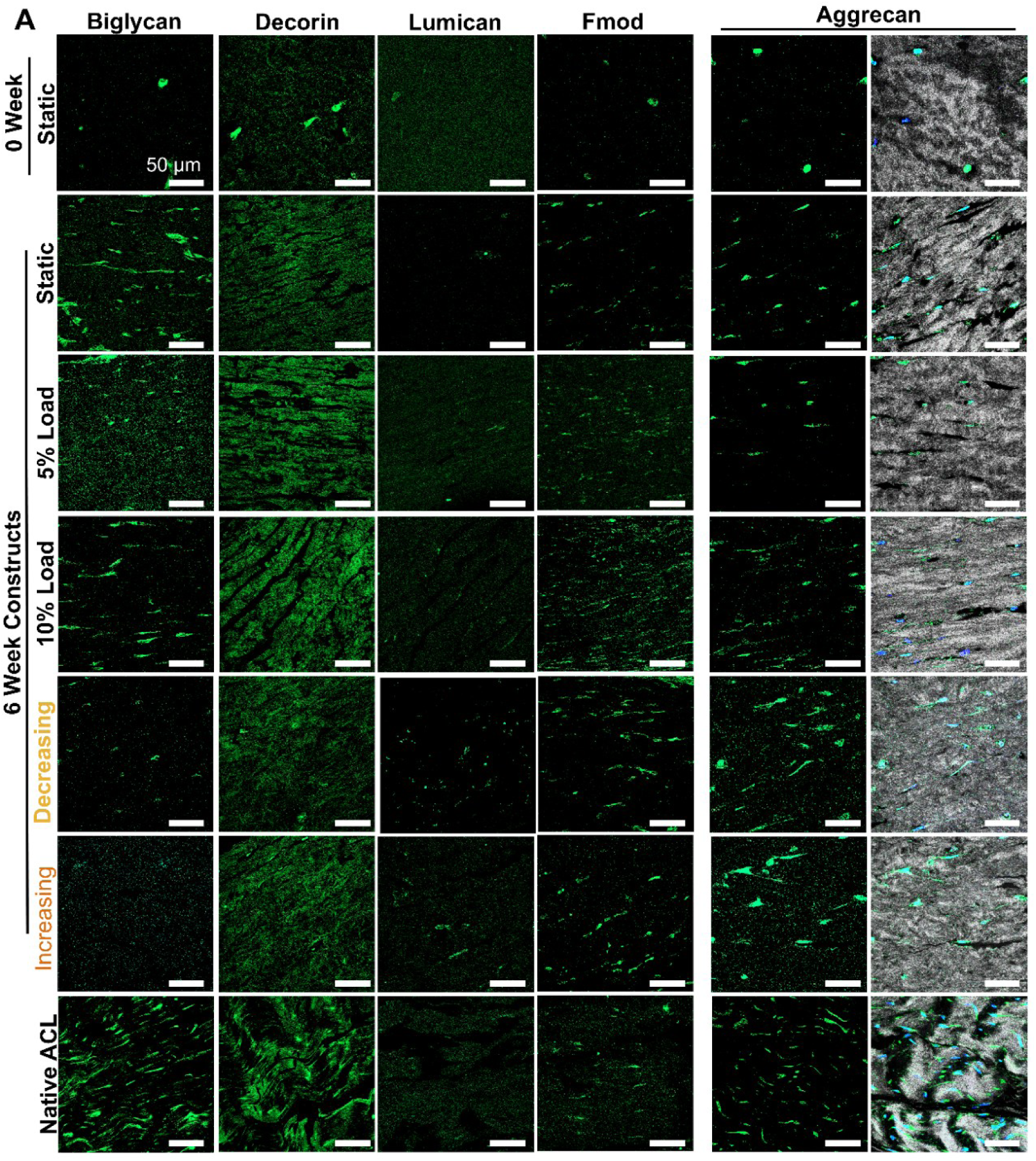
Stepped load drives variable accumulation of proteoglycans including biglycan, decorin, lumican, fibromodulin, and aggrecan. Immunofluorescence imaging of constructs at 0 and 6 weeks compared to neonatal bovine ACL (N = 3, n = 6 images per construct). First 5 columns: FITC = respective proteoglycan, final column: grey = collagen acquired from confocal reflectance imaging, DAPI = nuclei, FITC = aggrecan, scale bar = 50 µm for all. See Supplemental Figure 5 for representative negatives.

Quantification of these images revealed that all groups accumulated levels of decorin, lumican, fibromodulin, and aggrecan by 6 weeks that were not significantly different from neonatal bovine ACL, but none reached native levels of biglycan (**Figure 9**). Further, while static and steady load constructs had a significant increase in biglycan by 6 weeks, stepped load constructs did not. Interestingly, each loading condition drove different increases in SLRPs. Steady 5% load significantly increased decorin over all other groups to match native concentrations by 6 weeks. Decreasing load significantly increased lumican accumulation over all other groups and 10% steady load significantly increased fibromodulin over all other loading groups. Finally, both stepped loads accumulated more aggrecan than other treatments, but only decreasing load constructs had significantly increased aggrecan compared to other groups at 6 weeks.

**Figure 9.**
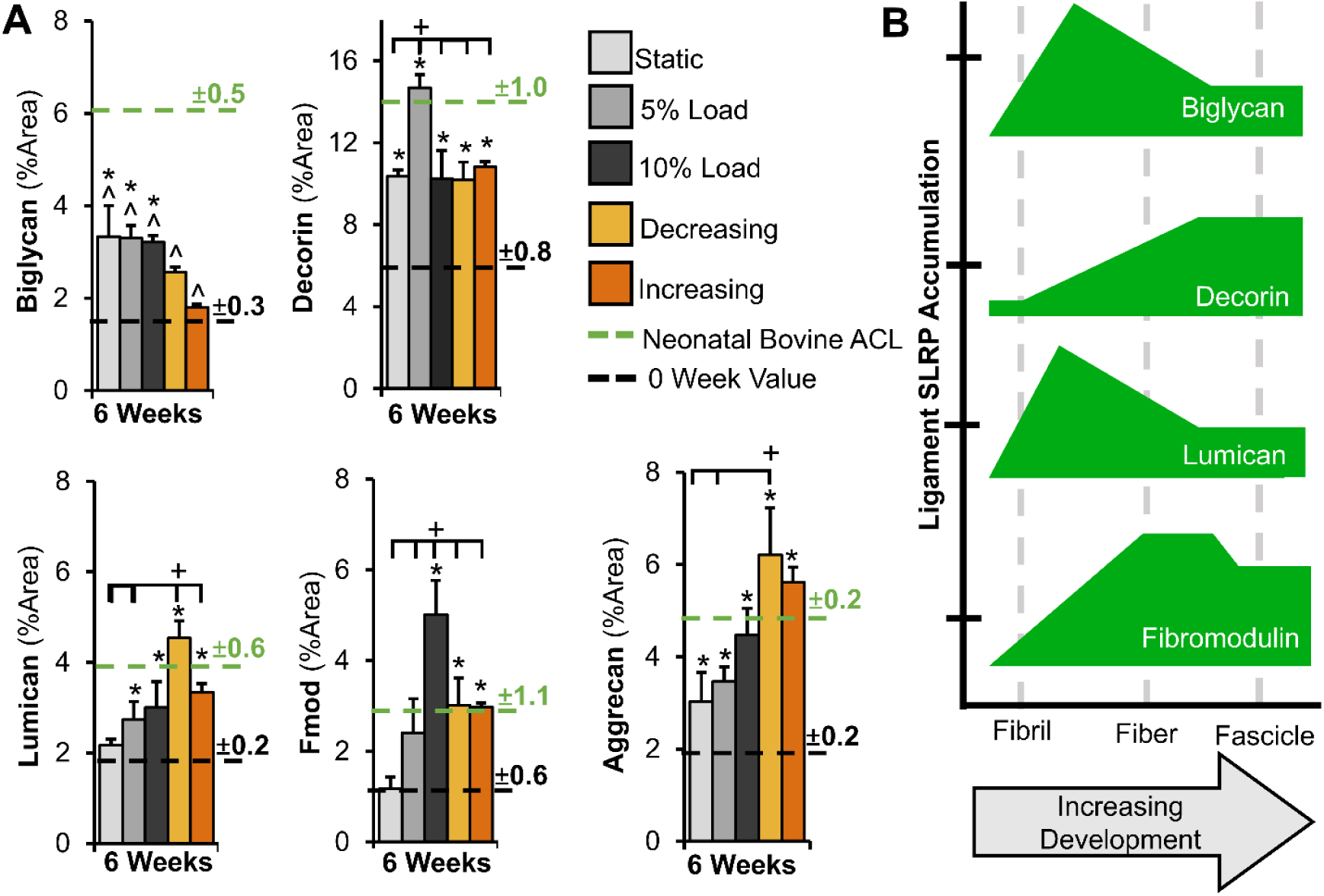
While Steady load constructs accumulated proteoglycans associated with later stages of hierarchical fiber formation, decreasing load constructs accumulated more lumican and aggrecan, associated with earlier stages of development and injury. A) Percent area measurements of positive immunofluorescent staining of biglycan, decorin, lumican, fibromodulin, and aggrecan in 0 week, 6 week, and neonatal bovine ACL tissues. Measurements from 6 images per sample were averaged to determine construct values (N = 3 constructs or native ACLs per treatment). Data shown as mean ± S.E.M. Significance compared to *0 week static, +bracket group, and ^neonatal bovine ACL (p < 0.05). B) Timeline schematic showing the relative expression of SLRPs throughout the hierarchical collagen maturation of ACL tissue (Based on literature findings [9, 65, 66, 103, 104]).

## 4. Discussion

The objective of this study was to explore whether a stepped cyclic load, that changed in strain magnitude as collagen fibers developed, further improved maturation in our engineered ligaments. Contrary to our hypothesis, neither decreasing nor increasing strain led to further maturation over steady load constructs. Previously, we found that steady 10% intermittent cyclic strain drove early improvements in maturation when cells were in unorganized gels and 5% strain was more beneficial later in culture once cells were anchored on aligned fibers [39], suggesting a decreasing stepped strain as collagen fibers form in our system may drive further maturation. However, here the decreasing load, with 10% strain early in culture and 5% strain later, did not lead to synergistic improvements in tissue maturation. Further, decreasing load constructs appeared to have a more injurious response to changes in strain, including swelling and elevated GAGs [1,9,74,75]. Additionally, the increasing load, which increased from 5% strain early in culture to 10% strain later, mirroring the increase in strain needed during development and healing *in vivo* [1,8], did not produce synergistic improvements over steady load constructs. In fact, both stepped load groups, whether decreasing or increasing, had significantly reduced hierarchical collagen organization, tensile properties, and composition compared to steady load and in some cases, static controls.

This lack of synergistic improvements in the stepped load constructs may be due to a disruption in tensional homeostasis. It is well established tenocytes and ligament fibroblasts form aligned collagen fibrils largely through cellular contraction forces, with cells remodeling the matrix in-order to achieve a tensional homeostasis [76–79]. Tensional homeostasis is an internal tensional load that cells attempt to maintain via contraction and remodeling of the extracellular matrix [78,79]. Mechanical stimulation, including cyclic load, which challenges this tensional homeostasis [77,78,80] has been shown to promote an anabolic response in cells and lead to increased tissue maturation [1,81,82]. However, disruptions in this tensional homeostasis, whether underload or overloading of the cell, often result in a catabolic response, which only shifts to anabolism once cells modify their environment to restore tensional homeostasis and sense the new load [77,79]. Thus, by suddenly changing the strain by 2-3% every 2 weeks, whether increasing or decreasing the strain, we might be disrupting cellular tensional homeostasis enough to cause cells to shift into a catabolic state, resulting in repeated remodeling and a reduction in tissue organization, strength, and composition. Further, while it is well established in engineered tissues that mechanical stimulation drives this repeated cycle of catabolism followed by anabolism and this repeated remodeling is what leads to increased tissue maturation[47,61,83], the 2 weeks of loading at a constant strain in this study may not be enough time for cells to re-establish their tensional homeostasis and drive substantial maturation.

More specifically, increasing and decreasing load constructs had reduced collagen organization compared to steady load constructs at the fibril, fiber and fascicle length-scale. In fact, both stepped loads had significantly reduced alignment at the fibril and fiber level and significantly reduced fiber diameters. This outcome is likely due to the continual turnover of collagen as cells attempt to remodel their surroundings in response to a change in load [77], as stepped applied strains changed every 2 weeks. However, stepped load constructs did have increased fibril diameters and a larger distribution of fibril diameters compared to static and 5% steady load. This may suggest that stepped strain drives improvements in fibril maturation, however the larger diameters in decreasing strain constructs may also be due to laxity in the tissue and subsequent lateral compression of the fibrils to restore tensional homeostasis, as has been previously observed in rat tail tendons [79,80].

In addition to disruption of fiber and fascicle maturation, stepped cyclic load further appeared to disrupt improvements in early crimp formation (**Figure 3 and 4**). Crimp is important to proper ligament function; however it is still unknown how it is formed [7,76,77]. This study provides valuable insight into the formation and disruption of crimp as fibers and fascicles form. All loaded groups appeared to develop crimp as early as 2 weeks, and while steady load constructs appeared to have continual improvements in uniformity and size of crimp with time in culture, stepped load constructs appeared to have smaller, more disorganized crimp formations by 6 weeks (**Figure 4A**). This result reflects previous work *in vivo* that found crimp length shifts with unloading, decreasing in length and becoming more irregular [80,84]. However, overloading and increased strains in ACL graphs has also been reported to drive increased crimp length in the first year of healing, but shift to shorter crimp length than native tissue 2-10 years post operation [85]. In increasing strain constructs we did not observe crimp lengthening, but instead smaller, more disorganized crimp. This may further suggest that the steps in strain from 5% to 7% to 10% throughout culture may be triggering a catabolic response, leading to remodeling of the tissue and thus more of an unloading effect in cells, rather than overload.

Along with reduced collagen organization, stepped load constructs also had reduced improvements in collagen accumulation with time in culture. Previously, with steady load, we found cells differentially regulated collagen accumulation depending on degree of organization and magnitude of load, with cyclic load first producing a catabolic response early in culture when cells were in unorganized gels, which later shifted to an anabolic response once cells were anchored on aligned fibrils and fibers [39]. However, here we found that after the strain changed at 2 weeks in stepped load constructs there was no further improvement in collagen concentration. In fact, increasing load constructs had a significant decrease in collagen concentration by 6 weeks. This reduction in collagen content and reduced organization in increasing load constructs reflects previous work in rat patellar tendon that found overload by high cyclic strain at 6 N reduces collagen concentration and organization [57]. A limitation to this study is that we did not directly measure collagen production or degradation, only bulk content. Future work should measure collagen discarded by cells into the media, as well as gene expression and matrix metalloproteinase production to better understand the changes in collagen production and degradation between loading conditions.

Interestingly, while decreasing load constructs did not significantly improve collagen accumulation, they did have a general increase in collagen concentration between 4 and 6 weeks to a final concentration not significantly different from neonatal bovine ACL. From 4 to 6 weeks, decreasing load constructs were being stimulated with 5% strain and the increase in collagen concentration during this period, mirrors the significant increases in collagen concentration found in 5% steady load constructs during this same period. This result may indicate that 5% strain is beneficial for collagen production when cells are anchored on aligned fibers. Traditionally, engineered tissues and injury recovery therapies are cyclically loaded at or below 5% strain, as it is the end range of motion for many tendons and ligaments *in vivo* [23,86,87], and this strain is reported to be optimal for tendon and ligament engineering [30,32–36].

Along with significantly reduced collagen accumulation, stepped load constructs also had significantly reduced LOX activity at 6 weeks compared to steady load groups, suggesting reduced collagen crosslinking. LOX driven crosslinking is paramount to tissue strength and previously it has been shown increased crosslinking reduces fibril size as it pulls collagen molecules tightly together [88,89]. Interestingly, decreasing load constructs had reduced LOX activity, increased fibril diameter, and reduced tensile strength, collectively suggesting reduced crosslinking. Further, It has been reported that LOX activity has a temporal component, and generally increases later in maturation as fascicles start to form [59,90,91]. At 6 weeks, steady load constructs increased in LOX activity, but stepped load constructs had a significant decrease in LOX activity compared to 4 weeks. This phenomenon could be due to a difference in maturation [59,90,91], a change in strain rate, or injury response [92,93].

Mirroring the lack of improvements in organization, collagen concentration, and LOX activity, tensile mechanical properties of stepped load constructs did not significantly improve over static clamped cultures in the elastic or toe region, and in the case of elastic modulus and UTS, performed significantly worse than clamped constructs. While steady load constructs had significant increases in tensile properties, reaching or surpassing immature native elastic modulus (1-3 MPa [94]) by 6 weeks, stepped load constructs did not reach or approach immature native tensile properties [94–96]. Collagen organization, concentration, and LOX crosslinking are all established to significantly affect the tensile strength of tendons and ligaments [8,88,97–100]. Thus, the reduced tensile strength in stepped load constructs correlates with their reduced organization, collagen concentration, and LOX activity. However, we did not directly measure LOX crosslinks. Future studies should evaluate the prevalence of divalent and trivalent LOX crosslinks to further investigate the correlation between these factors.

As mentioned previously, GAGs, and in particular, SLRPs serve an important role in fibrillogenesis [73,101,102]. Generally, SLRPs, including biglycan, decorin, lumican, and fibromodulin, are glycosylated proteins that interact with collagen fibrils, finely tuning and regulating fibrillogenesis [9,65,66,71]. Biglycan and lumican are present earlier in development, during fibril formation and early fiber assembly, but are later replaced by decorin and fibromodulin, respectively, as fibers bundle into larger fascicles (**Figure 9B**, based on literature findings [9,65,66,103,104]) Larger proteoglycans, like aggrecan also accumulate during development, serving a role in fiber spacing, load distribution, and water retention in response to compressive forces [9,65,69,70,105]. Aggrecan is critical for musculoskeletal tissue function, but an excess of this proteoglycan in tendons and ligaments is indicative of degeneration and injury [71,72].

Injuries in tendons and ligaments often result in increased GAG accumulation, which is theorized in part to be due to cells responding to a loss in tension [70,75]. In this study, static and steady load constructs had an increase in GAG accumulation over time in culture, leveling off at native concentrations, similar to previous studies [39]. However, decreasing load constructs had double the amount of GAG accumulation, significantly surpassing native levels. This could potentially suggest an injurious response to decreasing load, however which GAGs were being upregulated was not clear.

With immunohistochemistry analysis, we found steady and stepped load drove accumulation of different proteoglycans by 6 weeks. In particular, steady load drove increased accumulation of decorin and fibromodulin, depending on strain magnitude. As previously discussed, decorin and fibromodulin accumulate later in development, replacing biglycan and lumican, respectively, and are signs of increased maturation (**Figure 9B**). Higher levels of decorin in particular are associated with increased tensile mechanics [69], which is reflected with the increased tensile properties observed in steady load constructs. Fibromodulin also facilitates maturation as it allows for greater circumferential growth of fibers and facilitates crosslinking [73,106]. In general, high concentrations of biglycan, decorin, and fibromodulin are associated with lateral fibril growth and the spacing between fiber bundles, which modifies tensile properties [67,107–109]. On the other hand, stepped load constructs did not significantly increase accumulation of biglycan and decreasing load, in particular, had significant increases in lumican accumulation, suggesting reduced maturation. Increased lumican is generally associated with smaller, weaker fibers in ligament tissue [9,71,110], correlating well with the reduced collagen organization and tensile properties of decreasing load constructs.

Further, stepped load constructs accumulated more aggrecan than all other groups, with decreasing load constructs accumulating significantly more aggrecan than static and 5% steady load constructs. Increased aggrecan is associated with higher water retention and swelling [72,110], correlating with decreasing load constructs having a significantly higher percent wet weight than all other loaded groups at 4 and 6 weeks. Further, previous work has found that increased production of aggrecan leads to poorer fibril formation in meniscus tissue [18,30,31]. Our data correlates with this phenomenon in ligament, as stepped decreasing load constructs had significantly reduced hierarchical fiber formation and higher aggrecan accumulation. It has been established that changes in loading conditions, such as excessive strain or unloading, can elicit a maladaptive ligament fibroblast response, leading to collagen disorganization and a shift towards a more fibrocartilaginous tissue [114–118]. Fibrocartilaginous tissue generally accumulates higher levels of GAG than tendons and ligaments [9,69]. Therefore, the significantly increased GAG in decreasing load constructs at 6 weeks may indicate a potential injurious shift towards more fibrocartilaginous tissue [75,119,120].

Collectively, this study demonstrates that, contrary to our initial hypothesis, neither decreasing nor increasing stepped load, which changed as collagen organization improved, led to further improvements in hierarchical collagen organization or tissue mechanics. Previously, it has been shown that cells respond to changes in load with catabolism, which only shifts to anabolism once cells modify their environment to sense the new load [76,121]. We hypothesize stepped load did not further improve properties because the sudden 2-3% change in strain at 2 and 4 weeks was too large of a change in tensional homeostasis, for both increasing and decreasing, possibly inciting a catabolic response. However, a limitation of this study is we did not evaluate gene expression, molecules released to the media, or cellular contraction to further confirm these results. Future work should do traction force analysis, media, and gene expression analysis immediately following the change in strain and after the 2 weeks of constant strain to better confirm these results. If the change in strain is causing a catabolic response, smaller steps in strain, longer period between changes in strain to allow for more remodeling, or a more continuous ramped load or load-control regime may help drive further maturation in our system. Further, the intermittent loading regime used in this study (1 Hz, 1 hour on, 1 hour off, 1 hour on, 3x per week) is specifically tailored for optimal matrix turnover in unorganized gels, resulting in increased matrix accumulation [47,48,61,83,122], but changing any aspect, such as magnitude, frequency, duration, or rest period at any point in culture could change outcomes. Overall, this study provides new insight into how stepped intermittent cyclic load affects cell-driven hierarchical collagen fiber formation and how cells respond to changes in load over the course of culture. A better understanding of how mechanical cues differentially affect fiber formation in tissue-engineered tendons and ligaments will help improve loading protocols to drive collagen fiber maturation in engineered tissues and to develop better rehabilitation protocols to drive repair *in vivo*.

## 5. Conclusions

This study explored whether a stepped cyclic load that changes intensity as collagen fibers develop, further improves maturation. Contrary to our hypothesis, our findings indicate that neither decreasing nor increasing stepped cyclic strain, which changed as hierarchical fibers formed, led to further maturation. Instead, both stepped loads led to reduced hierarchical collagen organization compared to steady load constructs, and little to no improvements in tensile properties and composition. Previously, it has been shown that cells respond to changes in load with catabolism, which only shifts to anabolism once cells modify their environment to sense the new load [76,77]. These data may indicate that stepped cyclic load may lead to repeated breakdown and remodeling of collagen, which results in weaker, less organized construct than steady load which maintains cellular tensile homeostasis throughout culture [82,123]. In particular, decreasing load may lead to an injury response resulting in swelling, reduced organization, and shifted proteoglycan accumulation, including aggrecan. This study provides new insight into how cyclic loading affects cell-driven hierarchical fiber formation. A better understanding of how mechanical cues regulate fiber formation will help to better engineer replacements and develop better rehabilitation protocols to drive repair after injury

## Author Contribution

L.T. and J.L.P. conceived the project and wrote the manuscript. L.T. carried out experiments and analyzed the data. J.L.P. supervised the project and acquired funding. All authors edited the manuscript.

## Acknowledgements

The authors acknowledge the use of facilities within the Nanomaterials Characterization Core and the Virginia Commonwealth University Cancer Mouse Models Core Laboratory, supported in part, with funding from NIH-NCI Cancer Center Support Grant P30 CA016059.

## References

[1] M.T. Galloway, A.L. Lalley, J.T. Shearn, The Role of Mechanical Loading in Tendon Development, Maintenance, Injury, and Repair, J Bone Joint Surg Am 95 (2013) 1620–1628. 10.2106/JBJS.L.01004.

[2] K.E. Glattke, S.V. Tummala, A. Chhabra, Anterior Cruciate Ligament Reconstruction Recovery and Rehabilitation: A Systematic Review, JBJS 104 (2022) 739. 10.2106/JBJS.21.00688.

[3] K.L. Goh, A. Listrat, D. Béchet, Hierarchical Mechanics of Connective Tissues: Integrating Insights from Nano to Macroscopic Studies, Journal of Biomedical Nanotechnology 10 (2014) 2464–2507. 10.1166/jbn.2014.1960.

[4] G. Yang, B.B. Rothrauff, R.S. Tuan, Tendon and ligament regeneration and repair: Clinical relevance and developmental paradigm, Birth Defects Research Part C: Embryo Today: Reviews 99 (2013) 203–222. 10.1002/bdrc.21041.

[5] M. Pierantoni, I. Silva Barreto, M. Hammerman, V. Novak, A. Diaz, J. Engqvist, P. Eliasson, H. Isaksson, Multimodal and multiscale characterization reveals how tendon structure and mechanical response are altered by reduced loading, Acta Biomater 168 (2023) 264–276. 10.1016/j.actbio.2023.07.021.

[6] N. Karathanasopoulos, P. Angelikopoulos, C. Papadimitriou, P. Koumoutsakos, Bayesian identification of the tendon fascicle’s structural composition using finite element models for helical geometries, Computer Methods in Applied Mechanics and Engineering 313 (2017) 744–758. 10.1016/j.cma.2016.10.024.

[7] N. Karathanasopoulos, J. Ganghoffer, Investigating the Effect of Aging on the Viscosity of Tendon Fascicles and Fibers, Front. Bioeng. Biotechnol. 7 (2019). 10.3389/fbioe.2019.00107.

[8] B.K. Connizzo, S.M. Yannascoli, L.J. Soslowsky, Structure-function relationships of postnatal tendon development: a parallel to healing, Matrix Biol 32 (2013) 106–116. 10.1016/j.matbio.2013.01.007.

[9] S.C. Juneja, C. Veillette, Defects in Tendon, Ligament, and Enthesis in Response to Genetic Alterations in Key Proteoglycans and Glycoproteins: A Review, Arthritis 2013 (2013) 154812. 10.1155/2013/154812.

[10] R.B. Svensson, P. Hansen, T. Hassenkam, B.T. Haraldsson, P. Aagaard, V. Kovanen, M. Krogsgaard, M. Kjaer, S.P. Magnusson, Mechanical properties of human patellar tendon at the hierarchical levels of tendon and fibril, Journal of Applied Physiology 112 (2012) 419–426. 10.1152/japplphysiol.01172.2011.

[11] A. Gautieri, S. Vesentini, A. Redaelli, M.J. Buehler, Hierarchical Structure and Nanomechanics of Collagen Microfibrils from the Atomistic Scale Up, Nano Lett. 11 (2011) 757–766. 10.1021/nl103943u.

[12] Z. Chen, B. Zhou, X. Wang, G. Zhou, W. Zhang, B. Yi, W. Wang, W. Liu, Synergistic effects of mechanical stimulation and crimped topography to stimulate natural collagen development for tendon engineering, Acta Biomaterialia 145 (2022) 297–315. 10.1016/j.actbio.2022.04.026.

[13] T. Nau, A. Teuschl, Regeneration of the anterior cruciate ligament: Current strategies in tissue engineering, World J Orthop 6 (2015) 127–136. 10.5312/wjo.v6.i1.127.

[14] M.T. Rodrigues, R.L. Reis, M.E. Gomes, Engineering tendon and ligament tissues: present developments towards successful clinical products, Journal of Tissue Engineering and Regenerative Medicine 7 (2013) 673–686. 10.1002/term.1459.

[15] Y.J. No, M. Castilho, Y. Ramaswamy, H. Zreiqat, Role of Biomaterials and Controlled Architecture on Tendon/Ligament Repair and Regeneration, Advanced Materials 32 (2020) 1904511. 10.1002/adma.201904511.

[16] A.U. Khan, R. Aziz, M. Reen, W. Walker, P. Myers, The First Case of Bridge-Enhanced Anterior Cruciate Ligament (ACL) Repair (BEAR) Procedure in Mississippi, Cureus (2023). 10.7759/cureus.44218.

[17] M.M. Murray, B.C. Fleming, G.J. Badger, C. Freiberger, R. Henderson, S. Barnett, A. Kiapour, K. Ecklund, B. Proffen, N. Sant, D.E. Kramer, L.J. Micheli, Y.-M. Yen, Bridge-Enhanced Anterior Cruciate Ligament Repair Is Not Inferior to Autograft Anterior Cruciate Ligament Reconstruction at 2 Years: Results of a Prospective Randomized Clinical Trial, Am J Sports Med 48 (2020) 1305–1315. 10.1177/0363546520913532.

[18] V. Kandhari, T.D. Vieira, H. Ouanezar, C. Praz, N. Rosenstiel, C. Pioger, F. Franck, A. Saithna, B. Sonnery-Cottet, Clinical Outcomes of Arthroscopic Primary Anterior Cruciate Ligament Repair: A Systematic Review from the Scientific Anterior Cruciate Ligament Network International Study Group, Arthroscopy 36 (2020) 594–612. 10.1016/j.arthro.2019.09.021.

[19] E. Hing, D.K. Cherry, D.A. Woodwell, National Ambulatory Medical Care Survey: 2004 summary, Adv Data (2006) 1–33.

[20] S. Patel, J.-M. Caldwell, S.B. Doty, W.N. Levine, S. Rodeo, L.J. Soslowsky, S. Thomopoulos, H.H. Lu, Integrating soft and hard tissues via interface tissue engineering, Journal of Orthopaedic Research 36 (2018) 1069–1077. 10.1002/jor.23810.

[21] M.V. Paterno, S. Thomas, K.T. VanEtten, L.C. Schmitt, Confidence, ability to meet return to sport criteria, and second ACL injury risk associations after ACL-reconstruction, Journal of Orthopaedic Research 40 (2022) 182–190. 10.1002/jor.25071.

[22] Y. Wu, Y. Han, Y.S. Wong, J.Y.H. Fuh, Fibre-based scaffolding techniques for tendon tissue engineering, Journal of Tissue Engineering and Regenerative Medicine 12 (2018) 1798– 1821. 10.1002/term.2701.

[23] N.A. Dyment, J.G. Barrett, H.A. Awad, C.A. Bautista, A.J. Banes, D.L. Butler, A brief history of tendon and ligament bioreactors: Impact and future prospects, Journal of Orthopaedic Research 38 (2020) 2318–2330. 10.1002/jor.24784.

[24] C. Erisken, X. Zhang, K.L. Moffat, W.N. Levine, H.H. Lu, Scaffold Fiber Diameter Regulates Human Tendon Fibroblast Growth and Differentiation, Tissue Engineering Part A 19 (2013) 519–528. 10.1089/ten.tea.2012.0072.

[25] J.L. Puetzer, T. Ma, I. Sallent, A. Gelmi, M.M. Stevens, Driving Hierarchical Collagen Fiber Formation for Functional Tendon, Ligament, and Meniscus Replacement, Biomaterials 269 (2021) 120527. 10.1016/j.biomaterials.2020.120527.

[26] M.E. Brown, J.L. Puetzer, Driving native-like zonal enthesis formation in engineered ligaments using mechanical boundary conditions and β-tricalcium phosphate, Acta Biomaterialia 140 (2022) 700–716. 10.1016/j.actbio.2021.12.020.

[27] J. Ma, M.J. Smietana, T.Y. Kostrominova, E.M. Wojtys, L.M. Larkin, E.M. Arruda, Three-Dimensional Engineered Bone–Ligament–Bone Constructs for Anterior Cruciate Ligament Replacement, Tissue Engineering Part A 18 (2012) 103–116. 10.1089/ten.tea.2011.0231.

[28] B.M. Baker, R.P. Shah, A.H. Huang, R.L. Mauck, Dynamic Tensile Loading Improves the Functional Properties of Mesenchymal Stem Cell-Laden Nanofiber-Based Fibrocartilage, Tissue Engineering Part A 17 (2011) 1445–1455. 10.1089/ten.tea.2010.0535.

[29] L. Paschall, S. Carrozzi, E. Tabdanov, A. Dhawan, S.E. Szczesny, Cyclic loading induces anabolic gene expression in ACLs in a load-dependent and sex-specific manner, Journal of Orthopaedic Research 42 (2023) 267–276. 10.1002/jor.25677.

[30] K. Mubyana, D.T. Corr, Cyclic Uniaxial Tensile Strain Enhances the Mechanical Properties of Engineered, Scaffold-Free Tendon Fibers, Tissue Engineering Part A 24 (2018) 1808– 1817. 10.1089/ten.tea.2018.0028.

[31] W.K. Grier, R.A. Sun Han Chang, M.D. Ramsey, B.A.C. Harley, The influence of cyclic tensile strain on multi-compartment collagen-GAG scaffolds for tendon-bone junction repair, Connective Tissue Research 60 (2019) 530–543. 10.1080/03008207.2019.1601183.

[32] M.T.K. Bramson, S.K. Van Houten, D.T. Corr, Mechanobiology in Tendon, Ligament, and Skeletal Muscle Tissue Engineering, Journal of Biomechanical Engineering 143 (2021). 10.1115/1.4050035.

[33] N.R. Schiele, R.A. Koppes, D.B. Chrisey, D.T. Corr, Engineering Cellular Fibers for Musculoskeletal Soft Tissues Using Directed Self-Assembly, Tissue Engineering Part A 19 (2013) 1223–1232. 10.1089/ten.tea.2012.0321.

[34] E. Maeda, J.C. Shelton, D.L. Bader, D.A. Lee, Time dependence of cyclic tensile strain on collagen production in tendon fascicles, Biochemical and Biophysical Research Communications 362 (2007) 399–404. 10.1016/j.bbrc.2007.08.029.

[35] R.J. Crockett, M. Centrella, T.L. McCarthy, J. Grant Thomson, Effects of cyclic strain on rat tail tenocytes, Mol Biol Rep 37 (2010) 2629–2634. 10.1007/s11033-009-9788-8.

[36] J.A. Hannafin, S.P. Arnoczky, A. Hoonjan, P.A. Torzilli, Effect of stress deprivation and cyclic tensile loading on the material and morphologic properties of canine flexor digitorum profundus tendon: An in vitro study, Journal of Orthopaedic Research 13 (1995) 907–914. 10.1002/jor.1100130615.

[37] J.A. Hannafin, E.A. Attia, R. Henshaw, R.F. Warren, M.M. Bhargava, Effect of cyclic strain and plating matrix on cell proliferation and integrin expression by ligament fibroblasts, Journal of Orthopaedic Research 24 (2006) 149–158. 10.1002/jor.20018.

[38] J.L. Puetzer, L.J. Bonassar, High density type I collagen gels for tissue engineering of whole menisci, Acta Biomaterialia 9 (2013) 7787–7795. 10.1016/j.actbio.2013.05.002.

[39] L.D. Troop, J.L. Puetzer, Intermittent cyclic stretch of engineered ligaments drives hierarchical collagen fiber maturation in a dose- and organizational-dependent manner, Acta Biomaterialia 185 (2024) 296–311. 10.1016/j.actbio.2024.07.025.

[40] J.L. Chen, Z. Yin, W.L. Shen, X. Chen, B.C. Heng, X.H. Zou, H.W. Ouyang, Efficacy of hESC-MSCs in knitted silk-collagen scaffold for tendon tissue engineering and their roles, Biomaterials 31 (2010) 9438–9451. 10.1016/j.biomaterials.2010.08.011.

[41] M. Valdivia, F. Vega-Macaya, P. Olguín, Mechanical Control of Myotendinous Junction Formation and Tendon Differentiation during Development, Front. Cell Dev. Biol. 5 (2017). 10.3389/fcell.2017.00026.

[42] A.J. Banes, J. Gilbert, D. Taylor, O. Monbureau, A new vacuum-operated stress-providing instrument that applies static or variable duration cyclic tension or compression to cells in vitro, J Cell Sci 75 (1985) 35–42. 10.1242/jcs.75.1.35.

[43] J. Garvin, J. Qi, M. Maloney, A.J. Banes, Novel System for Engineering Bioartificial Tendons and Application of Mechanical Load, Tissue Engineering 9 (2003) 967–979. 10.1089/107632703322495619.

[44] K. Moe, T.E. Tay, J.C.H. Goh, H.W. Ouyang, S.L. Toh, Cyclic uniaxial strains on fibroblasts-seeded PLGA scaffolds for tissure engineering of ligaments, in: Third International Conference on Experimental Mechanics and Third Conference of the Asian Committee on Experimental Mechanics, SPIE, 2005: pp. 665–670. 10.1117/12.621764.

[45] G. Altman, R. Horan, I. Martin, Cell differentiation by mechanical stress, The FASEB Journal 16 (2001) 1–13. 10.1096/fj.01-0656fje.

[46] C. Androjna, R.K. Spragg, K.A. Derwin, Mechanical Conditioning of Cell-Seeded Small Intestine Submucosa: A Potential Tissue-Engineering Strategy for Tendon Repair, Tissue Engineering 13 (2007) 233–243. 10.1089/ten.2006.0050.

[47] G.D. Nicodemus, S.J. Bryant, Mechanical loading regimes affect the anabolic and catabolic activities by chondrocytes encapsulated in PEG hydrogels, Osteoarthritis and Cartilage 18 (2010) 126–137. 10.1016/j.joca.2009.08.005.

[48] M.E. Brown, J.L. Puetzer, Enthesis maturation in engineered ligaments is differentially driven by loads that mimic slow growth elongation and rapid cyclic muscle movement, Acta Biomaterialia 172 (2023) 106–122. 10.1016/j.actbio.2023.10.012.

[49] G.H. Altman, R.L. Horan, I. Martin, J. Farhadi, P.R.H. Stark, V. Volloch, J.C. Richmond, G. Vunjak-Novakovic, D.L. Kaplan, Cell differentiation by mechanical stress, The FASEB Journal 16 (2002) 1–13. 10.1096/fj.01-0656fje.

[50] J.M. Middendorf, M.E. Ita, B.A. Winkelstein, V. H. Barocas, Local tissue heterogeneity may modulate neuronal responses via altered axon strain fields: insights about innervated joint capsules from a computational model, Biomech Model Mechanobiol 20 (2021) 2269–2285. 10.1007/s10237-021-01506-9.

[51] G. McMahon, No Strain, No Gain? The Role of Strain and Load Magnitude in Human Tendon Responses and Adaptation to Loading, The Journal of Strength & Conditioning Research 36 (2022) 2950. 10.1519/JSC.0000000000004288.

[52] S. Bohm, F. Mersmann, A. Arampatzis, Human tendon adaptation in response to mechanical loading: a systematic review and meta-analysis of exercise intervention studies on healthy adults, Sports Med Open 1 (2015) 7. 10.1186/s40798-015-0009-9.

[53] A. Arampatzis, K. Karamanidis, K. Albracht, Adaptational responses of the human Achilles tendon by modulation of the applied cyclic strain magnitude, Journal of Experimental Biology 210 (2007) 2743–2753. 10.1242/jeb.003814.

[54] L. Pringels, J.L. Cook, E. Witvrouw, A. Burssens, L.V. Bossche, E. Wezenbeek, Exploring the role of intratendinous pressure in the pathogenesis of tendon pathology: a narrative review and conceptual framework, Br J Sports Med 57 (2023) 1042–1048. 10.1136/bjsports-2022-106066.

[55] G. DeMorat, P. Weinhold, T. Blackburn, S. Chudik, W. Garrett, Aggressive Quadriceps Loading Can Induce Noncontact Anterior Cruciate Ligament Injury, American Journal of Sports Medicine (2004). 10.1177/0363546503258928.

[56] P. Sbriccoli, M. Solomonow, B.-H. Zhou, Y. Lu, R. Sellards, Neuromuscular Response to Cyclic Loading of the Anterior Cruciate Ligament - Paola Sbriccoli, Moshe Solomonow, Bing-He Zhou, Yun Lu, Robert Sellards, 2005, The American Journal of Sports Medicine 33 (2005). 10.1177/0363546504268408.

[57] C. Hettrich, S. Gasinu, B. Beamer, M. Stasiak, A. Fox, P. Birmingham, O. Ying, D. Xiang-Hua, S. Rodeo, The Effect of Mechanical Load on Tendon-to-Bone Healing in a Rat Model - Carolyn M. Hettrich, Selom Gasinu, Brandon S. Beamer, Mark Stasiak, Alice Fox, Patrick Birmingham, Olivia Ying, Xiang-Hua Deng, Scott A. Rodeo, 2014, The American Journal of Sports Medicine 42 (2014). 10.1177/0363546514526.

[58] Investigación en Hemofilia y Fisioterapia, Efficacy of Load Control in Eccentric Exercises in the Increase of the Cross-sectional Area and Pain Threshold of the Patellar Tendon, and of the Strength of the Quadriceps in Volleyball Players. A Randomized Clinical Trial, (2020). https://clinicaltrials.gov/study/NCT03865862 (accessed November 4, 2025).

[59] M.E. Bates, L. Troop, M.E. Brown, J.L. Puetzer, Temporal application of lysyl oxidase during hierarchical collagen fiber formation differentially effects tissue mechanics, Acta Biomaterialia 160 (2023) 98–111. 10.1016/j.actbio.2023.02.024.

[60] V.L. Cross, Y. Zheng, N. Won Choi, S.S. Verbridge, B.A. Sutermaster, L.J. Bonassar, C. Fischbach, A.D. Stroock, Dense type I collagen matrices that support cellular remodeling and microfabrication for studies of tumor angiogenesis and vasculogenesis in vitro, Biomaterials 31 (2010) 8596–8607. 10.1016/j.biomaterials.2010.07.072.

[61] J.L. Puetzer, L.J. Bonassar, Physiologically Distributed Loading Patterns Drive the Formation of Zonally Organized Collagen Structures in Tissue-Engineered Meniscus, Tissue Engineering Part A 22 (2016) 907–916. 10.1089/ten.tea.2015.0519.

[62] J.L. Puetzer, E. Koo, L.J. Bonassar, Induction of fiber alignment and mechanical anisotropy in tissue engineered menisci with mechanical anchoring, Journal of Biomechanics 48 (2015) 1436–1443. 10.1016/j.jbiomech.2015.02.033.

[63] G. Kesava Reddy, C.S. Enwemeka, A simplified method for the analysis of hydroxyproline in biological tissues, Clinical Biochemistry 29 (1996) 225–229. 10.1016/0009-9120(96)00003-6.

[64] B.O. Enobakhare, D.L. Bader, D.A. Lee, Quantification of Sulfated Glycosaminoglycans in Chondrocyte/Alginate Cultures, by Use of 1,9-Dimethylmethylene Blue, Analytical Biochemistry 243 (1996) 189–191. 10.1006/abio.1996.0502.

[65] J.H. Yoon, J. Halper, Tendon proteoglycans: biochemistry and function, J Musculoskelet Neuronal Interact 5 (2005) 22–34.

[66] S. Chen, D.E. Birk, The regulatory roles of small leucine-rich proteoglycans in extracellular matrix assembly, FEBS J 280 (2013) 2120–2137. 10.1111/febs.12136.

[67] T. Douglas, S. Heinemann, S. Bierbaum, D. Scharnweber, H. Worch, Fibrillogenesis of Collagen Types I, II, and III with Small Leucine-Rich Proteoglycans Decorin and Biglycan, Biomacromolecules 7 (2006) 2388–2393. 10.1021/bm0603746.

[68] R.V. Iozzo, L. Schaefer, Proteoglycan form and function: A comprehensive nomenclature of proteoglycans, Matrix Biol 42 (2015) 11–55. 10.1016/j.matbio.2015.02.003.

[69] S.G. Lopez, L.J. Bonassar, The role of SLRPs and large aggregating proteoglycans in collagen fibrillogenesis, extracellular matrix assembly, and mechanical function of fibrocartilage, Connective Tissue Research 63 (2022) 269–286. 10.1080/03008207.2021.1903887.

[70] S. Mannion, A. Mtintsilana, M. Posthumus, W. Van Der Merwe, H. Hobbs, M. Collins, A.V. September, Genes encoding proteoglycans are associated with the risk of anterior cruciate ligament ruptures, Br J Sports Med 48 (2014) 1640–1646. 10.1136/bjsports-2013-093201.

[71] S. Ghatak, E. Maytin, J. Mack, V. Hascall, I. Atanelishvili, R. Rodriguez, R. Markwald, SunitiMisra, Roles of Proteoglycans and Glycosaminoglycans in Wound Healing and Fibrosis, International Journal of Cell Biol 2015 (2015). 10.1155/2015/834893.

[72] K.G. Vogel, J.D. Sandy, G. Pogány, J.R. Robbins, Aggrecan in bovine tendon, Matrix Biology 14 (1994) 171–179. 10.1016/0945-053X(94)90006-X.

[73] S. Kalamajski, A. Oldberg, The role of small leucine-rich proteoglycans in collagen fibrillogenesis, Matrix Biol 29 (2010) 248–253. 10.1016/j.matbio.2010.01.001.

[74] A.N. Corps, A.H.N. Robinson, T. Movin, M.L. Costa, B.L. Hazleman, G.P. Riley, Increased expression of aggrecan and biglycan mRNA in Achilles tendinopathy, Rheumatology (Oxford) 45 (2006) 291–294. 10.1093/rheumatology/kei152.

[75] J.E. Ackerman, K.T. Best, S.N. Muscat, A.E. Loiselle, Metabolic Regulation of Tendon Inflammation and Healing Following Injury, Curr Rheumatol Rep 23 (2021) 15. 10.1007/s11926-021-00981-4.

[76] M. Egerbacher, S.P. Arnoczky, O. Caballero, M. Lavagnino, K.L. Gardner, Loss of Homeostatic Tension Induces Apoptosis in Tendon Cells: An In Vitro Study, Clinical Orthopaedics and Related Research® 466 (2008) 1562. 10.1007/s11999-008-0274-8.

[77] M. Lavagnino, S.P. Arnoczky, In vitro alterations in cytoskeletal tensional homeostasis control gene expression in tendon cells, Journal of Orthopaedic Research 23 (2005) 1211– 1218. 10.1016/j.orthres.2005.04.001.

[78] K. Gardner, M. Lavagnino, M. Egerbacher, S.P. Arnoczky, Re-establishment of cytoskeletal tensional homeostasis in lax tendons occurs through an actin-mediated cellular contraction of the extracellular matrix, J Orthop Res 30 (2012) 1695–1701. 10.1002/jor.22131.

[79] S.P. Arnoczky, M. Lavagnino, M. Egerbacher, O. Caballero, K. Gardner, M.A. Shender, Loss of homeostatic strain alters mechanostat “set point” of tendon cells in vitro, Clin Orthop Relat Res 466 (2008) 1583–1591. 10.1007/s11999-008-0264-x.

[80] M. Lavagnino, A.E. Brooks, A.N. Oslapas, K.L. Gardner, S.P. Arnoczky, Crimp length decreases in lax tendons due to cytoskeletal tension, but is restored with tensional homeostasis, Journal of Orthopaedic Research 35 (2017) 573–579. 10.1002/jor.23489.

[81] M.L. Killian, L. Cavinatto, L.M. Galatz, S. Thomopoulos, The role of mechanobiology in tendon healing, J Shoulder Elbow Surg 21 (2012) 228–237. 10.1016/j.jse.2011.11.002.

[82] D.R. Henshaw, E. Attia, M. Bhargava, J.A. Hannafin, Canine ACL fibroblast integrin expression and cell alignment in response to cyclic tensile strain in three-dimensional collagen gels, Journal of Orthopaedic Research 24 (2006) 481–490. 10.1002/jor.20050.

[83] J.J. Ballyns, L.J. Bonassar, Dynamic compressive loading of image-guided tissue engineered meniscal constructs, Journal of Biomechanics 44 (2011) 509–516. 10.1016/j.jbiomech.2010.09.017.

[84] C.-H. Chen, X. Liu, M.-L. Yeh, M.-H. Huang, Q. Zhai, W.R. Lowe, D.M. Lintner, Z.-P. Luo, Pathological Changes of Human Ligament After Complete Mechanical Unloading, American Journal of Physical Medicine & Rehabilitation 86 (2007) 282. 10.1097/PHM.0b013e31803215dc.

[85] H.O. Mayr, A. Stoehr, K.T. Herberger, F. Haasters, A. Bernstein, H. Schmal, W.C. Prall, Histomorphological Alterations of Human Anterior Cruciate Ligament Grafts During Mid-Term and Long-Term Remodeling, Orthopaedic Surgery 13 (2021) 314–320. 10.1111/os.12835.

[86] S.E. Szczesny, D.T. Corr, Tendon cell and tissue culture: Perspectives and recommendations, Journal of Orthopaedic Research 41 (2023) 2093–2104. 10.1002/jor.25532.

[87] H. Delport, L. Labey, R. De Corte, B. Innocenti, J. Vander Sloten, J. Bellemans, Collateral ligament strains during knee joint laxity evaluation before and after TKA, Clinical Biomechanics 28 (2013) 777–782. 10.1016/j.clinbiomech.2013.06.006.

[88] E.A. Makris, R.F. MacBarb, D.J. Responte, J.C. Hu, K.A. Athanasiou, A copper sulfate and hydroxylysine treatment regimen for enhancing collagen cross-linking and biomechanical properties in engineered neocartilage, The FASEB Journal 27 (2013) 2421–2430. 10.1096/fj.12-224030.

[89] A.L. Cronlund, B.D. Smith, H.M. Kagan, Binding of Lysyl Oxidase to Fibrils of Type I Collagen, Connective Tissue Research 14 (1985) 109–119. 10.3109/03008208509015017.

[90] J.E. Marturano, J.F. Xylas, G.V. Sridharan, I. Georgakoudi, C.K. Kuo, Lysyl oxidase-mediated collagen crosslinks may be assessed as markers of functional properties of tendon tissue formation, Acta Biomaterialia 10 (2014) 1370–1379. 10.1016/j.actbio.2013.11.024.

[91] X.S. Pan, J. Li, E.B. Brown, C.K. Kuo, Embryo movements regulate tendon mechanical property development, Philosophical Transactions of the Royal Society B: Biological Sciences 373 (2018) 20170325. 10.1098/rstb.2017.0325.

[92] J. Xie, W. Huang, J. Jiang, Y. Zhang, Y. Xu, C. Xu, L. Yang, P.C.Y. Chen, K.L.P. Sung, Differential expressions of lysyl oxidase family in ACL and MCL fibroblasts after mechanical injury, Injury 44 (2013) 893–900. 10.1016/j.injury.2012.08.046.

[93] L. Cai, S. An, J. Liao, W. Yang, X. Zhou, K.P. Sung, J. Xie, Influences of Tumor Necrosis Factor–α on Lysyl Oxidases and Matrix Metalloproteinases of Injured Anterior Cruciate Ligament and Medial Collateral Ligament Fibroblasts, J Knee Surg 30 (2017) 78–87. 10.1055/s-0036-1581135.

[94] S.V. Eleswarapu, D.J. Responte, K.A. Athanasiou, Tensile Properties, Collagen Content, and Crosslinks in Connective Tissues of the Immature Knee Joint, PLOS ONE 6 (2011) e26178. 10.1371/journal.pone.0026178.

[95] S.G. McLean, K.F. Mallett, E.M. Arruda, Deconstructing the Anterior Cruciate Ligament: What We Know and Do Not Know About Function, Material Properties, and Injury Mechanics, Journal of Biomechanical Engineering 137 (2015). 10.1115/1.4029278.

[96] H. Fujie, Mechanical Properties and Biomechanical Function of the ACL, in: M. Ochi, K. Shino, K. Yasuda, M. Kurosaka (Eds.), ACL Injury and Its Treatment, Springer Japan, Tokyo, 2016: pp. 69–77. 10.1007/978-4-431-55858-3_6.

[97] E.A. Makris, D.J. Responte, N.K. Paschos, J.C. Hu, K.A. Athanasiou, Developing functional musculoskeletal tissues through hypoxia and lysyl oxidase-induced collagen cross-linking, Proceedings of the National Academy of Sciences 111 (2014) E4832–E4841. 10.1073/pnas.1414271111.

[98] M.J. Buehler, Nature designs tough collagen: Explaining the nanostructure of collagen fibrils, Proceedings of the National Academy of Sciences 103 (2006) 12285–12290. 10.1073/pnas.0603216103.

[99] C. Frank, D. McDonald, J. Wilson, D. Eyre, N. Shrive, Rabbit medial collateral ligament scar weakness is associated with decreased collagen pyridinoline crosslink density, Journal of Orthopaedic Research 13 (1995) 157–165. 10.1002/jor.1100130203.

[100] J.E. Marturano, J.D. Arena, Z.A. Schiller, I. Georgakoudi, C.K. Kuo, Characterization of mechanical and biochemical properties of developing embryonic tendon, Proceedings of the National Academy of Sciences 110 (2013) 6370–6375. 10.1073/pnas.1300135110.

[101] C.C. Banos, A.H. Thomas, C.K. Kuo, Collagen fibrillogenesis in tendon development: current models and regulation of fibril assembly, Birth Defects Res C Embryo Today 84 (2008) 228–244. 10.1002/bdrc.20130.

[102] R.V. Iozzo, The family of the small leucine-rich proteoglycans: key regulators of matrix assembly and cellular growth, Crit Rev Biochem Mol Biol 32 (1997) 141–174. 10.3109/10409239709108551.

[103] Y. Ezura, S. Chakravarti, Å. Oldberg, I. Chervoneva, D. Birk, Differential Expression of Lumican and Fibromodulin Regulate Collagen Fibrillogenesis in Developing Mouse Tendons | Journal of Cell Biology | Rockefeller University Press, Journal of Cell Biology 151 (2000) 779–788. 10.1083/jcb.151.4.779.

[104] H.L. Ansorge, S. Adams, D.E. Birk, L.J. Soslowsky, Mechanical, Compositional, and Structural Properties of the Post-natal Mouse Achilles Tendon, Ann Biomed Eng 39 (2011) 1904–1913. 10.1007/s10439-011-0299-0.

[105] S.G. Rees, C.M. Dent, B. Caterson, Metabolism of proteoglycans in tendon, Scandinavian Journal of Medicine & Science in Sports 19 (2009) 470–478. 10.1111/j.1600-0838.2009.00938.x.

[106] L. Svensson, A. Aszódi, F.P. Reinholt, R. Fässler, D. Heinegård, Å. Oldberg, Fibromodulin-null Mice Have Abnormal Collagen Fibrils, Tissue Organization, and Altered Lumican Deposition in Tendon*, Journal of Biological Chemistry 274 (1999) 9636–9647. 10.1074/jbc.274.14.9636.

[107] K.G. Danielson, H. Baribault, D.F. Holmes, H. Graham, K.E. Kadler, R.V. Iozzo, Targeted Disruption of Decorin Leads to Abnormal Collagen Fibril Morphology and Skin Fragility, J Cell Biol 136 (1997) 729–743. 10.1083/jcb.136.3.729.

[108] C.C. Reed, R.V. Iozzo, The role of decorin in collagen fibrillogenesis and skin homeostasis, Glycoconj J 19 (2002) 249–255. 10.1023/A:1025383913444.

[109] G. Zhang, Y. Ezura, I. Chervoneva, P.S. Robinson, D.P. Beason, E.T. Carine, L.J. Soslowsky, R.V. Iozzo, D.E. Birk, Decorin regulates assembly of collagen fibrils and acquisition of biomechanical properties during tendon development, Journal of Cellular Biochemistry 98 (2006) 1436–1449. 10.1002/jcb.20776.

[110] R.V. Iozzo, Matrix Proteoglycans: From Molecular Design to Cellular Function, Annual Review of Biochemistry 67 (1998) 609–652. 10.1146/annurev.biochem.67.1.609.

[111] S.G. Lopez, H.R. Moura, E. Chow, J.C.-H. Kuo, M.J. Paszek, L.J. Bonassar, Recombinant Small Leucine-Rich Proteoglycans Modulate Fiber Structure and Mechanical Properties of Collagen Gels, ACS Biomater. Sci. Eng. (2025). 10.1021/acsbiomaterials.5c00732.

[112] S.G. Lopez, J. Kim, L.A. Estroff, L.J. Bonassar, Removal of GAGs Regulates Mechanical Properties, Collagen Fiber Formation, and Alignment in Tissue Engineered Meniscus, ACS Biomater. Sci. Eng. 9 (2023) 1608–1619. 10.1021/acsbiomaterials.3c00136.

[113] S.G. Lopez, L.A. Estroff, L.J. Bonassar, siRNA Treatment Enhances Collagen Fiber Formation in Tissue-Engineered Meniscus via Transient Inhibition of Aggrecan Production, Bioengineering 11 (2024) 1308. 10.3390/bioengineering11121308.

[114] M. Stańczak, M. Biały, M. Hagner-Derengowska, Ligament Cell Biology: Effect of Mechanical Loading, Cell Physiol Biochem 59 (2025) 252–295. 10.33594/000000773.

[115] P.P.Y. Lui, Stem cell technology for tendon regeneration: current status, challenges, and future research directions, SCCAA 8 (2015) 163–174. 10.2147/SCCAA.S60832.

[116] S.C. Fu, W. Wang, H.M. Pau, Y.P. Wong, K.M. Chan, C.G. Rolf, Increased Expression of Transforming Growth Factor-β1 in Patellar Tendinosis, Clinical Orthopaedics and Related Research® 400 (2002) 174.

[117] K. Legerlotz, G.P. Riley, H.R.C. Screen, GAG depletion increases the stress-relaxation response of tendon fascicles, but does not influence recovery, Acta Biomaterialia 9 (2013) 6860–6866. 10.1016/j.actbio.2013.02.028.

[118] N.L. Millar, M. Akbar, A.L. Campbell, J.H. Reilly, S.C. Kerr, M. McLean, M. Frleta-Gilchrist, U.G. Fazzi, W.J. Leach, B.P. Rooney, L.A.N. Crowe, G.A.C. Murrell, I.B. McInnes, IL-17A mediates inflammatory and tissue remodelling events in early human tendinopathy, Sci Rep 6 (2016) 27149. 10.1038/srep27149.

[119] J. Nyland, A. Huffstutler, J. Faridi, S. Sachdeva, M. Nyland, D. Caborn, Cruciate ligament healing and injury prevention in the age of regenerative medicine and technostress: homeostasis revisited, Knee Surg Sports Traumatol Arthrosc 28 (2020) 777–789. 10.1007/s00167-019-05458-7.

[120] H.-J. Jung, M.B. Fisher, S.L.-Y. Woo, Role of biomechanics in the understanding of normal, injured, and healing ligaments and tendons, BMC Sports Sci Med Rehabil 1 (2009) 9. 10.1186/1758-2555-1-9.

[121] M. Lavagnino, M.E. Wall, D. Little, A.J. Banes, F. Guilak, S.P. Arnoczky, Tendon mechanobiology: Current knowledge and future research opportunities, Journal of Orthopaedic Research 33 (2015) 813–822. 10.1002/jor.22871.

[122] J.D. Kisiday, J.H. Lee, P.N. Siparsky, D.D. Frisbie, C.R. Flannery, J.D. Sandy, A.J. Grodzinsky, Catabolic Responses of Chondrocyte-Seeded Peptide Hydrogel to Dynamic Compression, Ann Biomed Eng 37 (2009) 1368–1375. 10.1007/s10439-009-9699-9.

[123] M. Lavagnino, S.P. Arnoczky, M. Egerbacher, K.L. Gardner, M.E. Burns, Isolated fibrillar damage in tendons stimulates local collagenase mRNA expression and protein synthesis, Journal of Biomechanics 39 (2006) 2355–2362. 10.1016/j.jbiomech.2005.08.008.

