## Supplemental for "Stepped Cyclic Strain, that Increases or Decreases as Hierarchical Collagen Fibers Form, Does not Further Improve Maturation in Engineered Ligaments"

### Supplemental Figures:

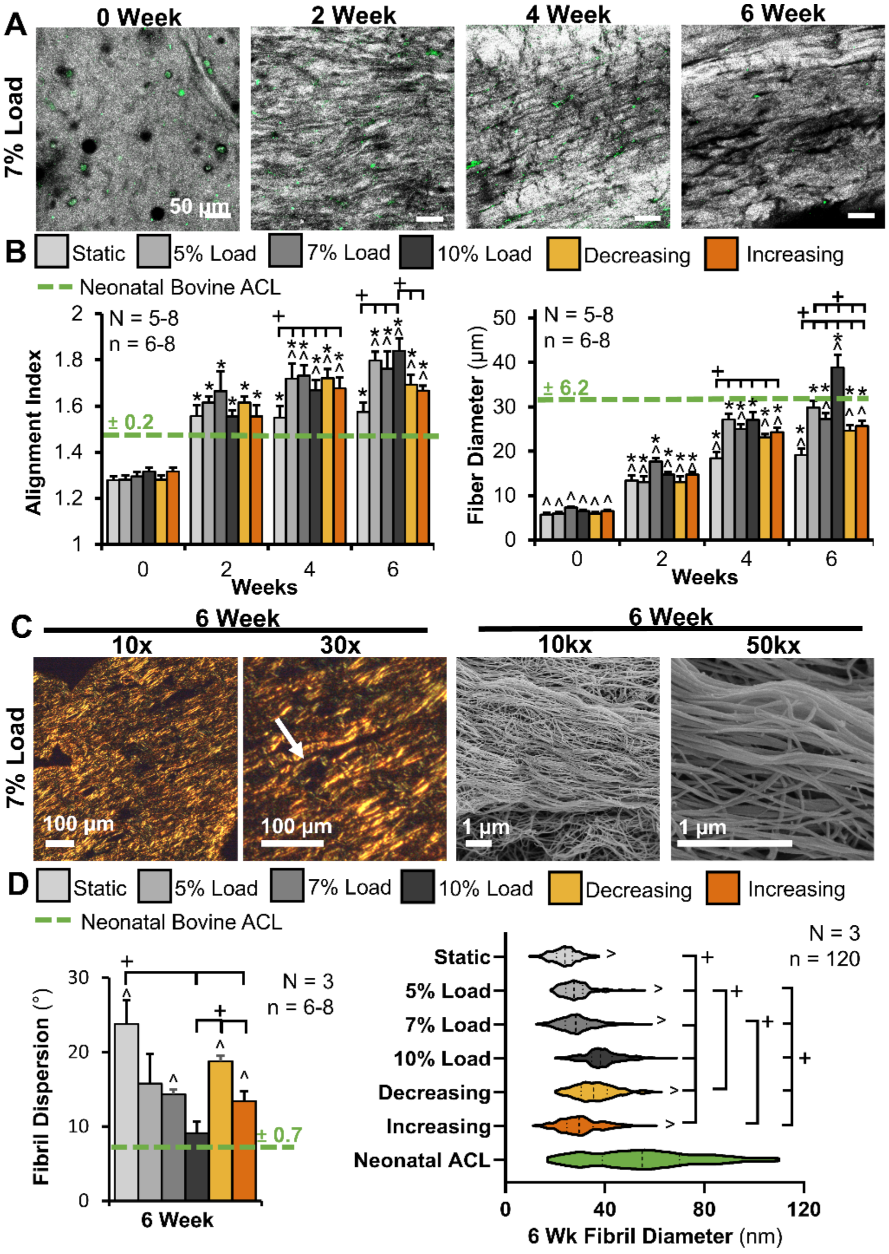


**Supplemental Figure 1:** Steady 7% load data largely remained between outcomes of 5% and 10% steady load. A) Confocal reflectance of 7% loaded constructs throughout culture. Grey = collagen, green = cellular auto-fluorescence, scale bar = 50 µm. B) Degree of collagen alignment (reported via alignment index where 1 is unorganized, 4.5 is perfectly aligned) and average collagen fiber diameter determined via a FFT-based image analysis. 6–8 confocal images per construct were averaged to determine construct values, with 5–8 constructs analyzed per time point and 4 native samples analyzed. C) Fascicle length-scale organization at 6 weeks evaluated by picrosirius red staining, imaged with polarized light (scale bar = 100 µm), and SEM images of 6-week constructs to assess fibril length-scale organization (scale bar = 1 µm). D) Analysis of SEM images to determine fibril dispersion (lower dispersion indicating increased alignment), and fibril diameter of 6-week constructs and neonatal bovine ACL. For dispersion, 6–8 images per construct were averaged to determine the construct average (N = 3 for constructs and native ACL). For fibril diameters, 20 fibrils per image for 6 images were measured and pooled (total 120 fibrils per construct) to determine average fibril diameter for each construct. All fibril diameters from $N = 3$ constructs were pooled (n = 360) to visualize the spread of fibril diameters at 6 weeks. Data shown as mean ± S.E.M. Significance compared to *0 week static, ^neonatal bovine ACL or +bracket group ($p < 0.05$)

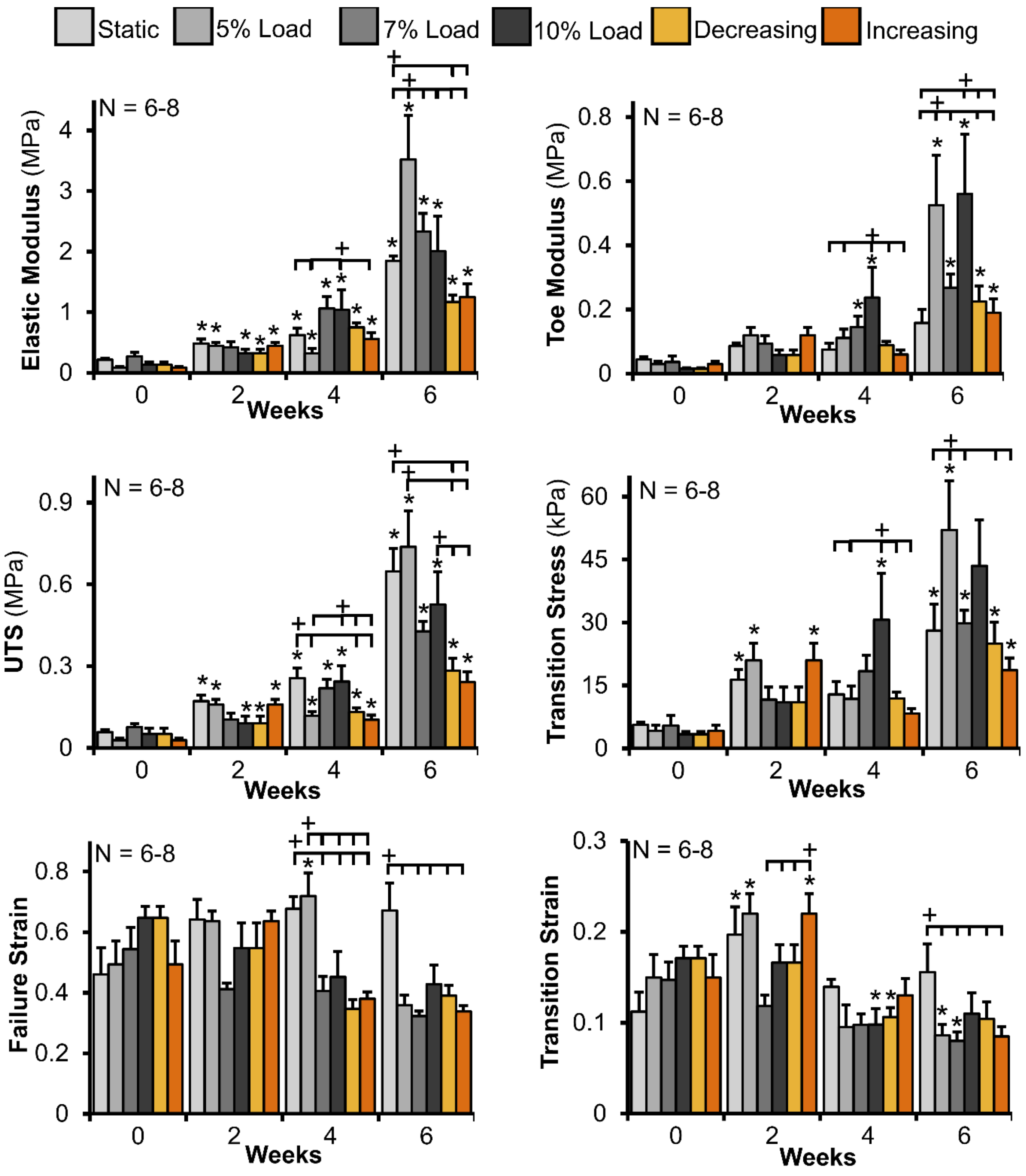


**Supplemental Figure 2:** Steady 7% load data largely remained between outcomes of 5% and 10% steady load for A) elastic properties (Elastic modulus, ultimate tensile strength (UTS), and failure strain) and B) toe-region properties (Toe modulus, transition stress, and transition strain) over 6 weeks of culture. *N* = 6–8. Data shown as mean ± S.E.M. Significance compared to *0 week static and +bracket group ($p < 0.05$).

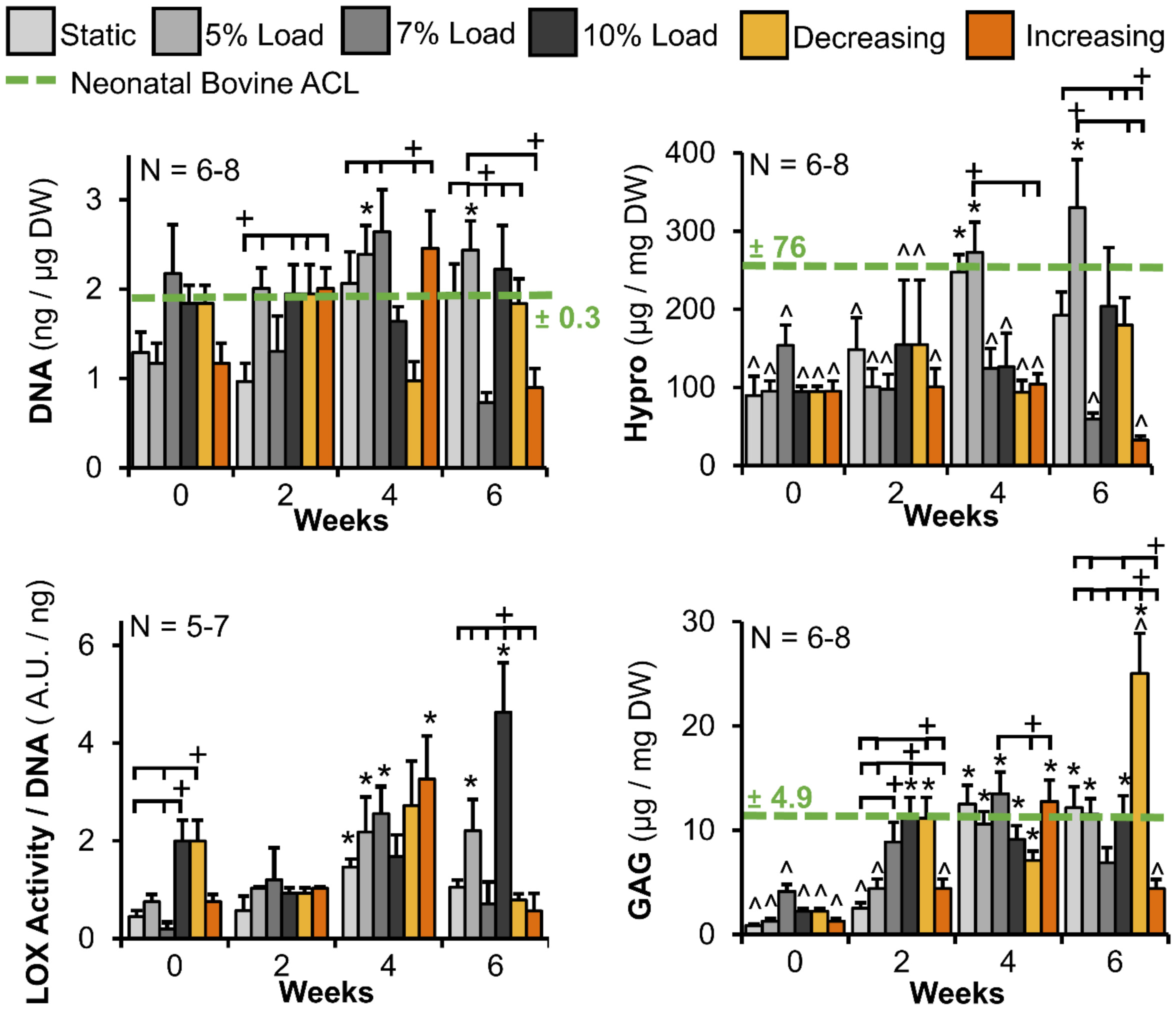


**Supplemental Figure 3:** DNA, collagen content (represented by hydroxyproline), and GAG, normalized to dry weight (N = 6-8), and LOX activity normalized to DNA (N = 5-7). Data shown as mean ± S.E.M. Significance compared to *0 week static, ^neonatal bovine ACL, and +bracket group ($p$ < 0.05).

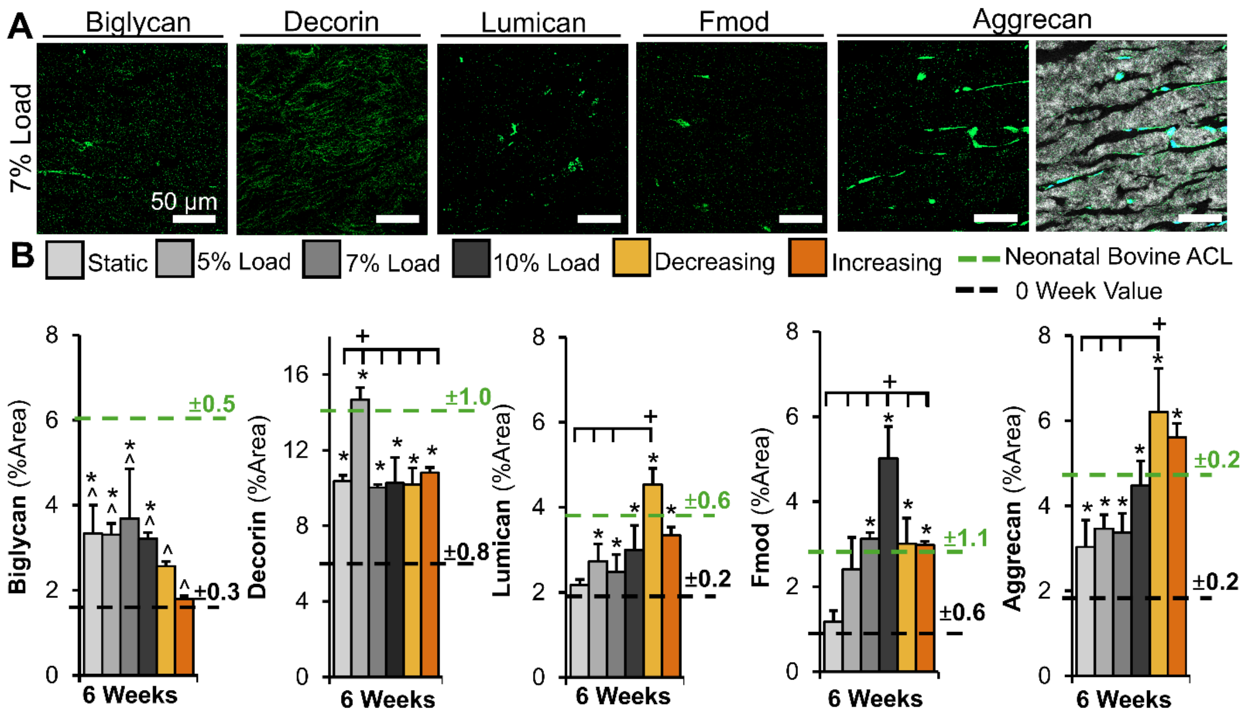


**Supplemental Figure 4:** Steady 7% cyclic load accumulated proteoglycans largely similar to 5% and 10% steady load. A) Representative IHC images of 7% steady load constructs stained for proteoglycans. First 5 columns: FITC = respective proteoglycan, final column: grey = collagen, DAPI = nuclei, FITC = aggrecan, scale bar = 50 μm for all. B) Percent area measurements of positive immunofluorescent straining of biglycan, decorin, lumican, fibromodulin, and aggrecan in 0 week, 6 week, and neonatal bovine ACL tissues. Measurements from 6 images per sample were averaged to determine construct values (N = 3 constructs or native ACLs per treatment). Data shown as mean ± S.E.M. Significance compared to *0 week static, +bracket group, and ^neonatal bovine ACL ($p$ < 0.05).

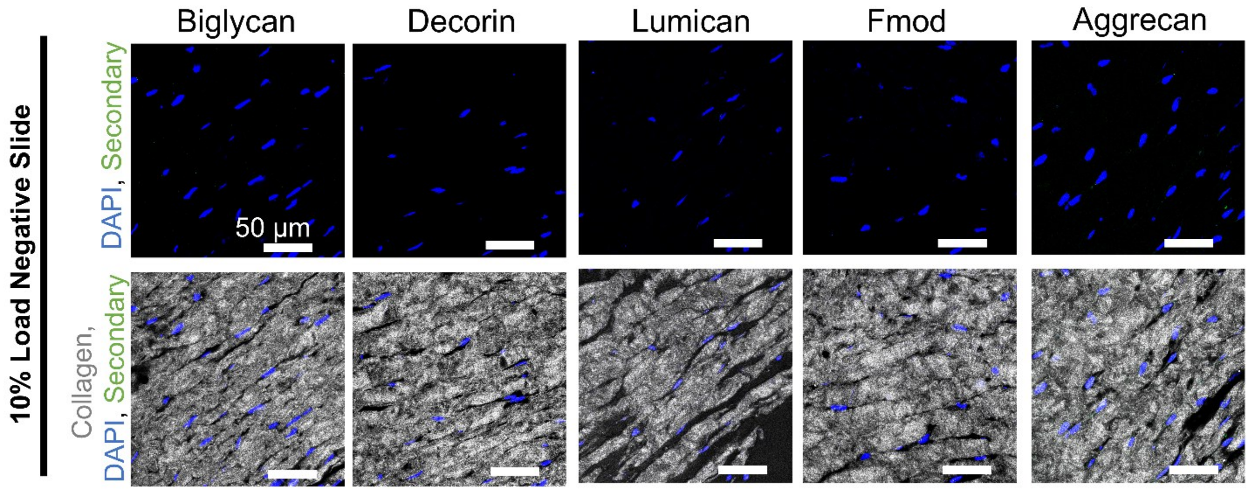


**Supplemental Figure 5:** Representative confocal reflectance images of negative immunofluorescence samples for proteoglycan localization. FITC = respective proteoglycan, grey = collagen acquired from confocal reflectance imaging, DAPI = nuclei, scale bar = 50 μm.

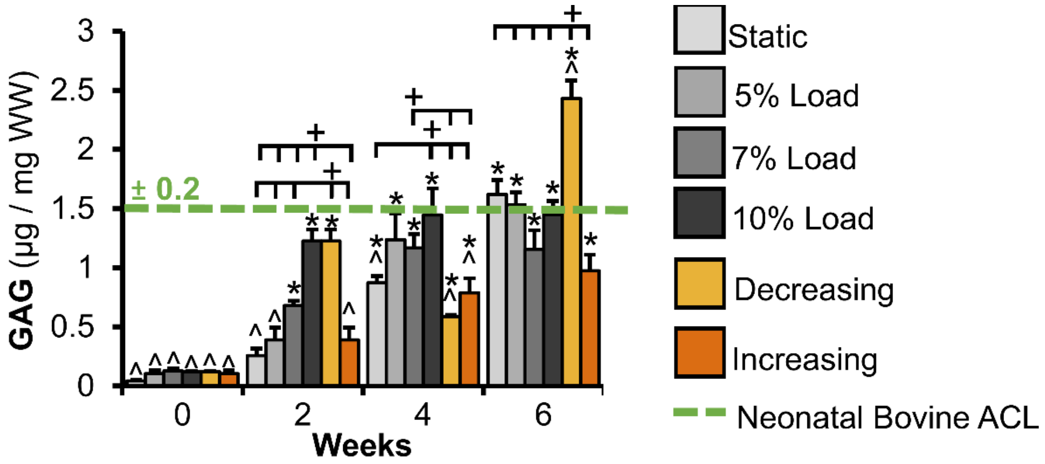


**Supplemental Figure 6:** Construct GAG accumulation normalized to wet weight (WW, N = 6-8). Data shown as mean ± S.E.M. Significance compared to *0 week static, ^neonatal bovine ACL, and +bracket group ($p < 0.05$).
